# Imaging the Transmembrane and Transendothelial Sodium Gradients in Gliomas

**DOI:** 10.1101/2020.08.26.268839

**Authors:** Muhammad H. Khan, John J. Walsh, Jelena M. Mihailović, Sandeep K. Mishra, Daniel Coman, Fahmeed Hyder

## Abstract

High sodium (Na^+^) in extracellular (Na^+^_e_) and blood (Na^+^_b_) compartments and low Na^+^ in intracellular milieu (Na^+^_i_) produce strong transmembrane (ΔNa^+^_mem_) and weak transendothelial (ΔNa^+^_end_) gradients respectively, which reflect cell membrane potential (*V_m_*) and blood-brain barrier (BBB) integrity. We developed a sodium (^23^Na) magnetic resonance spectroscopic imaging (MRSI) method using an intravenously-administered paramagnetic contrast agent to measure ΔNa^+^_mem_ and ΔNa^+^_end_. *In vitro* ^23^Na-MRSI established that the ^23^Na signal is strongly shifted by the agent compared to biological factors. *In vivo* ^23^Na-MRSI showed Na^+^_i_ remained unshifted and Na^+^_b_ was more shifted than Na^+^_e_, and these together created weakened ΔNa^+^_mem_ and enhanced ΔNa^+^_end_ in rat gliomas. Specifically, RG2 and U87 tumors maintained weakened ΔNa^+^_mem_ (i.e., depolarized *V_m_*) implying an aggressive state for proliferation, and RG2 tumors displayed elevated ΔNa^+^_end_ suggesting altered BBB integrity. ^23^Na-MRSI will allow explorations of perturbed Na^+^ homeostasis *in vivo* for the tumor neurovascular unit.

## INTRODUCTION

Sodium (Na^+^) concentration is normally low intracellularly (~10 mM) and high in blood and extracellular spaces (~150 mM)(Bean, 2007; Cheng et al., 2013; Ennis et al., 1996), producing a strong transmembrane Na^+^ gradient (ΔNa^+^_mem_~140 mM) and a weak transendothelial Na^+^ gradient (ΔNa^+^_end_ ~0 mM). The ΔNa^+^_mem_ is coupled to the cell membrane potential (*V_m_*), nerve signaling(Bean, 2007), muscle activity(Juel, 1986) and osmoregulation(Stock et al., 2002), while the ΔNa^+^_end_ impacts bicarbonate and proton transport between extracellular and blood spaces(Boron, 2004; Ennis et al., 1996; Green et al., 1986; Hladky & Barrand, 2016) and signifies blood-brain barrier (BBB) integrity(Shah & Kimberly, 2016; Stokum et al., 2015).

The sodium-potassium pump transports Na^+^ against its electrochemical gradient by consuming adenosine triphosphate generated through oxidative phosphorylation(Cheng et al., 2013). In glioblastoma (GBM), glycolysis is upregulated in relation to oxidative phosphorylation even with sufficient oxygen(DeBerardinis & Chandel, 2016). This aerobic glycolysis generates excessive amounts of hydrogen ions and lactate intracellularly, which are extruded into the extracellular milieu, lowering the pH of the tumor microenvironment(Hyder & Manjura Hoque, 2017). Since both the cell membrane and BBB regulate the ionic composition of the extracellular fluid(Bean, 2007; Ennis et al., 1996), we posited that maintaining ΔNa^+^_mem_ and ΔNa^+^_end_ becomes unsustainable in the tumor neurovascular unit. Since cancer is the second-leading cause of death globally(Koene et al., 2016), measuring [Na^+^] across different compartments *in vivo* has potential to become an invaluable biomarker.

Hyperpolarized *V_m_* corresponds to quiescent cell cycle stages (*G_0_* phase), and depolarized *V_m_* to proliferative/replicative stages (*M* phase)(Cone, 1970; JOHNSTONE, 1959; Yang & Brackenbury, 2013). Therefore, ΔNa^+^_mem_ is a biomarker for tumorigenicity and tumor aggressiveness. Determining [Na^+^] in the extracellular milieu usually involves inserting microelectrodes through the skull and reading voltage differences across cellular compartments(Petersen, 2017). Besides issues of accurate microelectrode positioning and tissue penetration, such invasive techniques challenge human translation.

Angiogenesis is a crucial part of tumor growth(Folkman, 2006). Unlike normal tissues, the immature tumor vasculature exhibits saccular formations, hyperbranching, and twisted patterns that cause the BBB to be leaky. Prior cancer research avoided measuring [Na^+^] in blood presumably due to microhemorrhage concerns from ruptured blood vessels with microelectrodes. But given the gamut of anti-angiogenic therapies for GBM(Batchelor et al., 2014), measuring ΔNa^+^_end_ non-invasively is desirable.

Nuclear magnetic resonance (NMR) non-invasively detects the isotope sodium-23 (^23^Na), a spin-3/2 quadrupolar nucleus. ^23^Na is 100% abundant and provides the second-strongest signal *in vivo*, next to hydrogen (^1^H) which is spin-1/2 and non-quadrupolar(Anderson et al., 1978). ^23^Na magnetic resonance imaging (MRI) has greatly impacted stroke and ischemia research(Hilal et al., 1983; Moseley et al., 1985), but can only reflect total sodium (Na^+^_T_)(Madelin, Lee, et al., 2014; Madelin, Poidevin, et al., 2015) because resonances from blood (Na^+^_b_), extracellular (Na^+^_e_), and intracellular (Na^+^_i_) compartments are difficult to separate. Thus, quantification of transmembrane (ΔNa^+^_mem_ = Na^+^_e_ - Na^+^_i_) and transendothelial (ΔNa^+^_end_ = Na^+^_b_ - Na^+^_e_) gradients has eluded ^23^Na-MRI. While detecting Na^+^_T_ is useful clinically, ΔNa^+^_end_ and ΔNa^+^_mem_ can reveal relevant information about BBB viability and cellular proliferative/oncogenic potential. ^23^Na-MRI methods based on diffusion, inversion recovery, and multiple quantum filtering (MQF) attempt to separate Na^+^_i_ and Na^+^_e_ signals and their volume fractions, but suffer from low sensitivity and necessitate large magnetic field gradients due to low gyromagnetic ratio (γ_Na_) and short longitudinal/transverse relaxation times (*T_1_/T_2_*) for ^23^Na. These ^23^Na-MRI methods need enhanced specificity for the Na^+^_i_ signal because they cannot fully suppress major contributions from Na^+^_b_ and Na^+^_e_, both of which dominate the Na^+^_T_ signal(Madelin, Babb, et al., 2015; Madelin, Kline, et al., 2014; Madelin, Lee, et al., 2014).

Another approach to separate Na^+^ signals *in vivo* involves intravenous administration of an exogenous paramagnetic but polyanionic contrast agent (paraCA^*n*-^). The paraCA^*n*-^ consists of a lanthanide(III) cation core bound to an anionic macrocyclic chelate(Chu et al., 1984; Chu et al., 1990). Since the paraCA^*n*-^ extravasates into extracellular space of most organs but not cells, only Na^+^_e_ and Na^+^_b_ will be attracted to the paraCA^*n*-^ and experience a shift in the ^23^Na resonance frequency to separate the ^23^Na magnetic resonance spectroscopic imaging (MRSI) signals between Na^+^_b_, Na^+^_e_ and Na^+^_i_. Proof-of-concept for this has been demonstrated *in situ* for the heart(Weidensteiner et al., 2002) and liver(Colet et al., 2005). Given the compromised BBB in tumors relative to healthy tissue, the ^23^Na-MRSI technique in conjunction with paraCA^*n*-^ is particularly efficacious in studying brain tumors.

The most effective paraCA^*n*-^ for compartmental ^23^Na separation is(Bansal et al., 1992) the thulium(III) cation (Tm^3+^) complexed with 1,4,7,10-tetraazacyclododecane-1,4,7,10-tetrakis(methylenephosphonate) (DOTP^8-^) to form TmDOTP^5-^ (**Figure 1(a)**). TmDOTP^5-^ has enjoyed many *in vivo* applications, both with ^1^H-NMR(Coman et al., 2016; Coman, Trubel, & Hyder, 2009; Coman, Trubel, Rycyna, et al., 2009; Huang et al., 2016) and ^23^Na-NMR(Colet et al., 2005; Ronen & Kim, 2001). Particularly, TmDOTP^5-^ has been infused intravenously to induce ^23^Na compartmental signal separation in healthy(Bansal et al., 1992; Winter et al., 1998) and tumor-bearing rats(Winter & Bansal, 2001). However, these studies looked at non-localized ^23^Na signals, which obfuscate the results due to the ubiquity of Na^+^ in other tissues. Our goal was to investigate ΔNa^+^_mem_ and ΔNa^+^_end_ in tumor and normal tissues in 3D using ^23^Na-MRSI with TmDOTP^5-^ at high spatial resolution.

**Figure 1.**
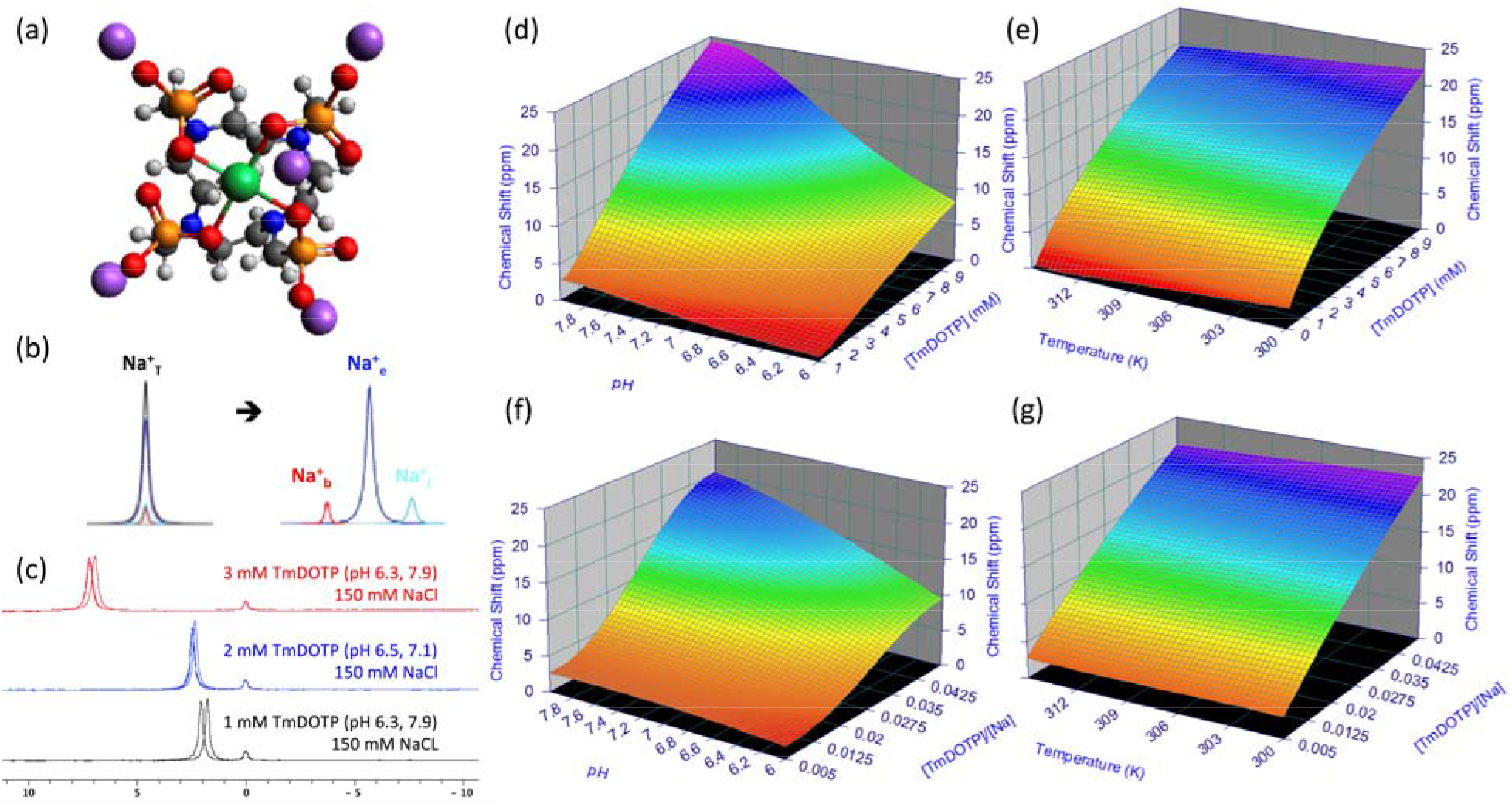
Shifting mechanism of the ^23^Na resonance. (a) Chemical structure of sodium thulium(III) 1,4,7,10-tetraazacyclododecane-1,4,7,10-tetrakis(methylenephosphonate) (Na_5_TmDOTP). The TmDOTP^5-^ complex consists of the Tm^3+^ ion chelated with DOTP^8-^. Each phosphonate-containing pendant arm on TmDOTP^5-^ has electron-donating groups on the oxygen atoms(red) to stabilize the Tm^3+^ conjugation with DOTP^8-^. The −5 charge simultaneously attracts five Na^+^ ions(purple), which experience a shift in the observed ^23^Na resonance that is dependent on [TmDOTP^5-^]. (b) *In vivo*, prior to TmDOTP^5-^ administration(left), the ^23^Na spectrum yields only a single peak representing the total sodium(Na^+^_T_) comprising blood(Na^+^_b_), extracellular(Na^+^_e_), and intracellular(Na^+^_i_) compartments. Following TmDOTP^5-^ administration(right), the peaks become spectroscopically separable based on [TmDOTP^5-^] in each compartment. Integrals of these peaks will be representative of [Na^+^] in each compartment. (c) A two-compartment coaxial cylinder tube setup was employed for *in vitro* observation of the chemical shift separation scheme **(Figure S1)**. The inner tube (smaller volume) was filled with 150 mM NaCl, while the outer tube (larger volume) was filled with the same solution in addition to various amounts of TmDOTP^5-^, each subject to different pH conditions. Thus, all ^23^Na spectra from this phantom setup displayed a small unshifted peak from the inner compartment and a larger shifted peak. The outer-to-inner volume ratio was 8.6, explaining the difference in sizes. Exemplary traces of ^23^Na spectra show that the shift is much more sensitive to [TmDOTP^5-^] (2.77 ppm/mM) than to variations in pH (0.25 ppm/pH unit) or temperature (0.03 ppm/°C). Plots (d) and (e) show that temperature, pH, and [TmDOTP^5-^] all contribute to variations of the ^23^Na chemical shift. However, these plots depict ranges of pH and temperature that are unlikely for *in vivo* settings (i.e., changes over 2 full pH units and temperature changes over 15 °C). Moreover, [Na^+^] *in vivo* (~150 mM in blood and extracellular space) is extremely high compared to [TmDOTP^5-^] (~2 mM in blood, ~1 mM in extracellular space). Therefore, variations in ^23^Na chemical shift are primarily dependent on [TmDOTP^5-^]/[Na^+^] thereby rendering (f) pH and (g) temperature dependencies negligible. Data points were fit to Chebyshev rational polynomials using TableCurve 3D.

*In vitro* studies established that the ^23^Na shift is more sensitive to [TmDOTP^5-^] than other biological factors such as changes in pH and/or temperature (**Figure 1(b-g); Supplementary: Theory**). Upon *in vivo* administration of TmDOTP^5-^, three peaks were observed, corresponding to Na^+^_b_, Na^+^_e_, and Na^+^_i_. Na^+^_b_ was shifted the most (2 ppm) while Na^+^_i_ remained unshifted. Our *in vivo* results, consistent with prior studies of tumor cells *in vitro*(Yang & Brackenbury, 2013), demonstrated a significantly weakened ΔNa^+^_mem_ and strengthened ΔNa^+^_end_ within tumor tissue relative to healthy tissue as consequences of elevated Na^+^_b_ and lowered Na^+^_e_, respectively. The ^23^Na vascular results showed similar patterns to traditional vascular imaging by ^1^H-based dynamic contrast-enhanced MRI (^1^H-DCE-MRI)(Sourbron & Buckley, 2013). We describe nuances of these novel measurements of disrupted Na^+^ homeostasis in cancer and their implications.

## RESULTS

### *In Vitro* Studies for Mechanistic Separation of ^23^Na Peaks

The goal of these studies was to separate the total Na^+^ signal (Na^+^_T_) into distinct signals for blood (Na^+^_b_), extracellular (Na^+^_e_), and intracellular (Na^+^_i_) pools (**Figure 1(b)**). The shifting mechanism induced by exogenous TmDOTP^5-^ and endogenous biological factors on the ^23^Na chemical shift *in vitro* is shown in **Figure 1(c-g)**. A two-compartment coaxial cylinder NMR tube setup *in vitro* was used to mimic Na^+^ in extracellular/intracellular pools *in vivo* (**Figure S1**). The inner (smaller) and outer (larger) compartments both contained 150 mM NaCl while the latter also contained TmDOTP^5-^ at various concentrations. The whole setup was subjected to several different pH and temperature conditions. Since the inner compartment lacked TmDOTP^5-^, all spectra exhibited a small, unshifted peak at 0 ppm. The larger peak was shifted downfield by TmDOTP^5-^, with the difference in peak integrals stemming from different compartment volumes (**Figure S1**). To demonstrate the feasibility of this approach to quantify Na^+^ signals from different compartments, the contents of the compartments were switched and the above measurements were repeated (**Figure S1**).

*In vitro* ^23^Na spectra revealed that the chemical shift is most sensitive to [TmDOTP^5-^], compared to pH and temperature (**Figure 1(c)**). The ^23^Na shiftability for TmDOTP^5-^ (*s*_[*paraCA^n−^*]_=2-77 ppm/mM; **Equation (3)** in **Supplementary: Theory**) was 11.1× larger than the shiftability for pH (*s_pH_*=0.25 ppm/pH unit) and 92.3× larger than the shiftability for temperature (*s_T_*=0.03 ppm/°C). This means that addition of 1.1 mM TmDOTP^5-^ would induce a ~3 ppm shift in the ^23^Na peak. Conversely, a maximal change of 0.4 in pH units, which is seen between normal and tumor tissues(Coman et al., 2016), would induce only a ~0.1 ppm shift. A similar shift by temperature would require a 3.3-°C change, which is unlikely *in vivo*. Based on the pH and temperature ranges observed *in vivo* (including tumors), the effect from TmDOTP^5-^ dominates the chemical shift (**Equation (2)** in **Supplementary: Theory**) by 95%. Therefore, [TmDOTP^5-^] is several orders of magnitude more sensitive in shifting the ^23^Na resonance than typical *in vivo* biological factors. Consequently, ^23^Na spectra displayed dependence mostly on [TmDOTP^5-^] (**Figures 1(d)-(e)**). However for *in vivo* scenarios the ranges shown for pH (2 full pH units) and temperature (15-°C interval) are overestimated, and where [Na^+^] far exceeds [TmDOTP^5-^] based on prior experiments(Coman, Trubel, Rycyna, et al., 2009). In blood and extracellular spaces, [Na^+^] is ~30-100× greater than [TmDOTP^5-^](Coman, Trubel, & Hyder, 2009). This suggests that the *relative amount* of TmDOTP^5-^ is the primary factor affecting ^23^Na chemical shift (**Equation (4)** in **Supplementary: Theory**).

### *In Vivo* Separation of ^23^Na Peaks Indicates Compartmentalized Na^+^ Pools

Interrogating individual voxels in the brain before and after TmDOTP^5-^ administration revealed clear ^23^Na signal separation, although to varying extents depending on the degree of TmDOTP^5-^ extravasation from blood to the extracellular space. ^23^Na-MRSI data overlaid on ^1^H-MRI anatomy of rat brains bearing U251 tumors showed spectra in tumor and healthy tissue voxels (**Figure 2**), with candidate voxels inside/outside the tumor before and after TmDOTP^5-^. Before TmDOTP^5-^ delivery, there was a single ^23^Na peak observed at 0 ppm corresponding to Na^+^_T_ both inside and outside the tumor. Upon TmDOTP^5-^ delivery, compartmental ^23^Na peak separation was achieved. Within the tumor, the compromised BBB permitted greater TmDOTP^5-^ extravasation and accumulation in the extracellular space, explicitly yielding three separate peaks emerging from the original single ^23^Na resonance. Each peak was associated with a compartment, with Na^+^_i_ being the unshifted peak (0 ppm) because TmDOTP^5-^ could not enter the intracellular compartment. The most-shifted peak (~2 ppm) was Na^+^_b_ because the blood compartment had the largest [TmDOTP^5-^]. This was corroborated by blood samples removed from the animal and observing a shifted peak at ~2 ppm in the ^23^Na NMR spectrum. The intermediate peak in the middle corresponded to the extracellular Na^+^_e_ resonance. The splitting was also evident outside of the tumor (i.e., in healthy tissue) where TmDOTP^5-^ extravasated to a much lesser extent compared to tumor tissue (**Figure 2**). The Na^+^_b_ peak was observed at ~2 ppm, whereas the Na^+^_i_ and Na^+^_e_ peaks were less discernible. The shifted bulk Na^+^_e_ peak confirmed that whatever degree of TmDOTP^5-^ extravasation occurred was sufficient to affect the extracellular ^23^Na signals, albeit less pronounced than tumoral Na^+^_e_. The unshifted Na^+^_i_ resonance was still at 0 ppm, but partially eclipsed by the bulk Na^+^_e_ peak. These same patterns inside/outside the tumor were observed throughout the brain (see **Figure S2** for voxels in the same rat).

**Figure 2.**
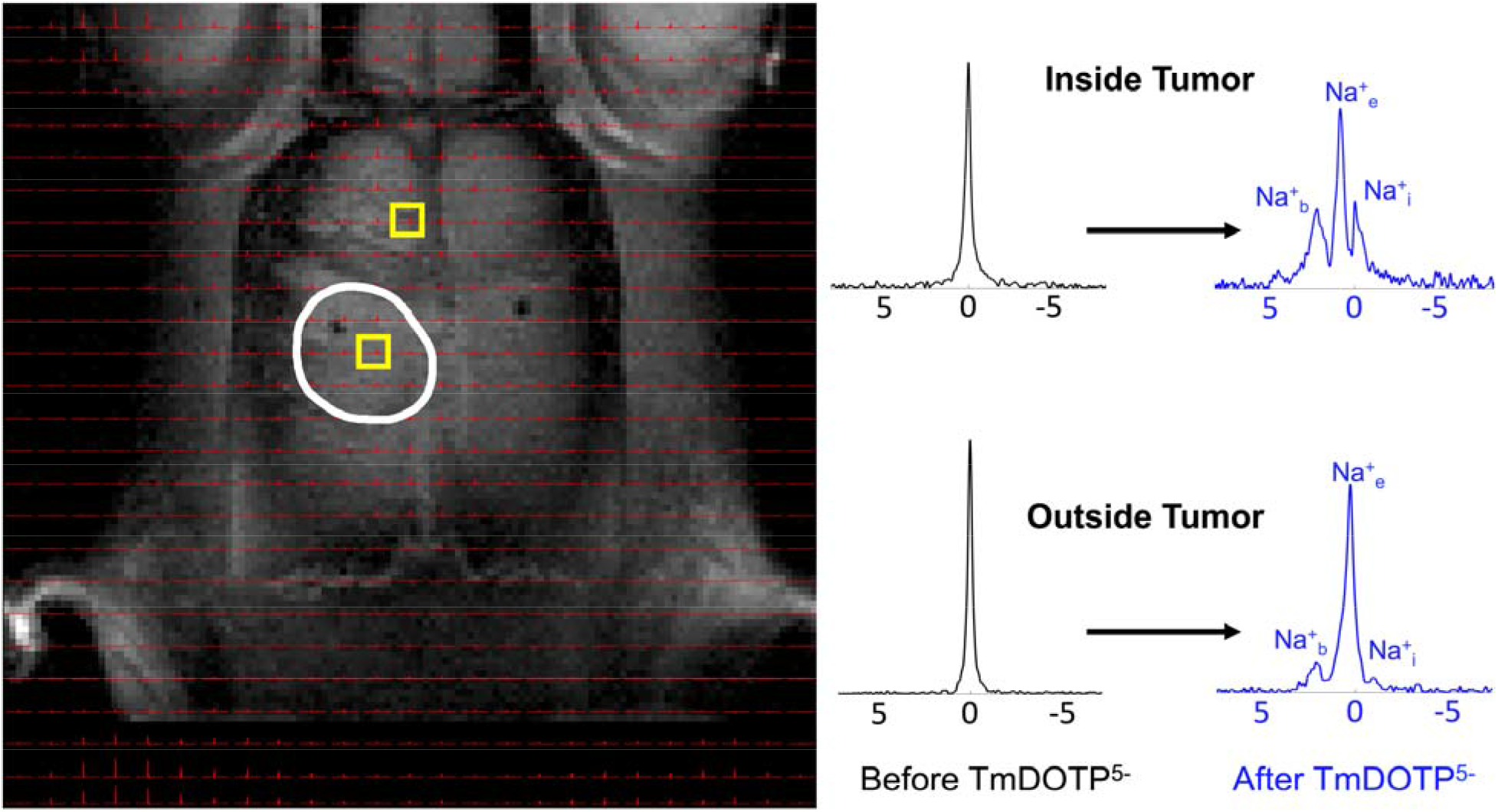
Demonstration of ^23^Na peak separation *in vivo* following TmDOTP^5-^ administration into a rat brain bearing a U251 tumor. ^1^H-MRI of an axial slice displaying the anatomical tumor boundary (white outline). The ^23^Na-MRSI is overlaid on top of the ^1^H-MRI. Candidate voxels inside and outside the tumor are indicated (yellow boxes). Before TmDOTP^5-^ delivery, a single ^23^Na peak was observed at 0 ppm, corresponding to total sodium (Na^+^_T_), both inside and outside the tumor (black spectra). Following TmDOTP^5-^ delivery, compartmental peak separation was achieved to varying extents throughout the brain (blue spectra). Within the tumor (top spectra), this separation was most pronounced due to a compromised blood-brain barrier (BBB), which permits substantial accumulation of TmDOTP^5-^ in the extracellular space. Outside of the tumor (bottom spectra), such a high degree of extravasation would not be possible, but some shifting is still observed. The TmDOTP^5-^ distribution in the brain warrants labeling the most shifted peak as blood sodium (Na^+^_b_), which occurred consistently around 2 ppm. The unshifted peak, which has no access to TmDOTP^5-^, is intracellular sodium (Na^+^_i_). The intermediate peak, therefore, is extracellular sodium (Na^+^_e_), which is shifted more inside the tumor than outside in healthy tissue. Similar spectroscopic patterns are observed throughout all voxels *in vivo*. See **Figure S2** for a slice below the present. All spectra were magnitude-corrected and line-broadened by 10 Hz.

**Figure 3** displays data from representative rats bearing (a) RG2 and (b) U87 tumors, with the array of ^23^Na-MRSI data overlaid on top of the ^1^H-MRI anatomy. The spectra from individual voxels placed throughout the brain confirmed only one ^23^Na peak prior to infusion (**Figure 3, black spectra**), corresponding to Na^+^_T_, but upon infusion the single peak separated into two additional peaks (**Figure 3, green spectra**).

**Figure 3.**
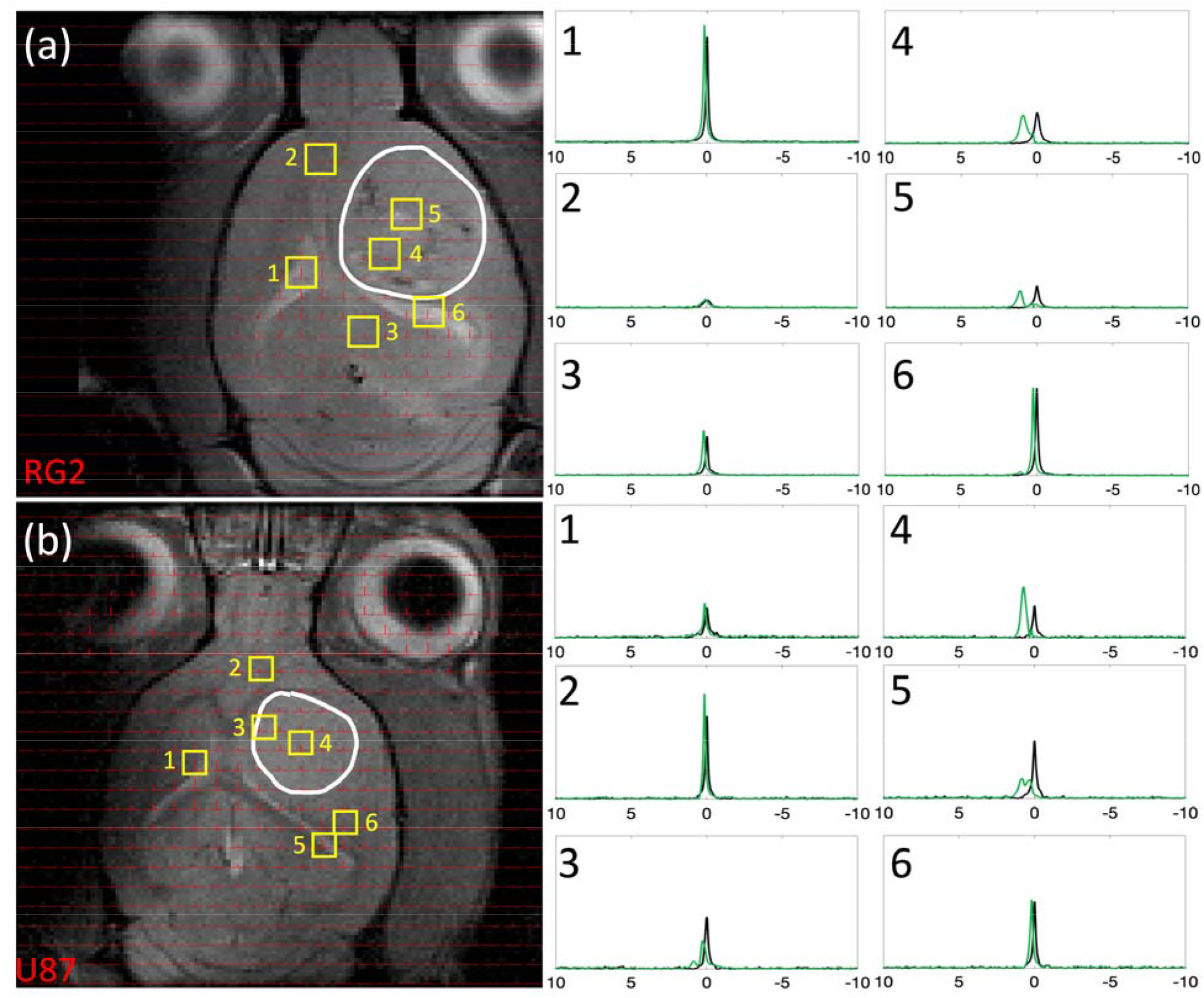
Comparison of ^23^Na peak separation in rats bearing RG2 and U87 tumors. For rats bearing an (a) RG2 and (b) U87 tumor, the tumor boundary is outlined in white, with voxels of interest indicated in yellow squares (with numbers), and spectra acquired before and after TmDOTP^5-^ delivery shown in black and green, respectively. Tumor voxels [(a) 4 and 5 in RG2 tumor, (b) 3 and 4 in U87 tumor] exhibited a fair amount of peak separation due to the leaky BBB. Na^+^_b_ shift was consistently around 2 ppm, and Na^+^_i_ shift was at 0 ppm, whereas Na^+^_e_ shift in the tumor was in the range 0.5-1 ppm. Voxels in the healthy tissue [(a) 2 and 3 in RG2 tumor, (b) 2 and 6 in U87 tumor] were slightly shifted in the positive direction, suggesting the paramagnetic effects of TmDOTP^5-^ reach the extracellular space even with limited extravasation. Ventricular voxels [(a) 1 and 6 in RG2 tumor, (b) 1 and 5 in U87 tumor] displayed a single unshifted Lorentzian peak before and a shifted Lorentzian peak after TmDOTP^5-^ injection. This is attributed to the dominant ^23^Na signal contribution in the ventricles coming from cerebrospinal fluid (CSF), which contains free (i.e., unbound) aqueous Na^+^. The position of the shifted ventricle peak coincided with the Na^+^_e_ peak position in other regions of the brain. This agrees with expectation because CSF is in physical contact with the extracellular space with free exchange of aqueous Na^+^ between the two compartments. Similar spectroscopic patterns are observed throughout all voxels *in vivo*. See **Figure S3** for several slices for each rat shown here. All spectra were magnitude-corrected and line-broadened by 10 Hz.

Prior to TmDOTP^5-^ infusion (**Figure 3, black spectra**) ventricular voxels [**Figure 3(a)**: voxels 1,6; **Figure 3(b)**: voxels 1,5] exhibited purely Lorentzian lineshapes characterized by a single *T*_2_, while those in the normal brain [**Figure 3(a)**: voxels 2,3; **Figure 3(b)**:2,6] and tumor [**Figure 3(a)**: voxels 4,5; **Figure 3(b)**: voxels 3,4] displayed super-Lorentzian lineshapes indicative of multiple *T*_2_ values. This is because the ventricles are comprised almost entirely of cerebrospinal fluid (CSF), in which all Na^+^ is aqueous, whereas some Na^+^ ions in tissue can be bound. These observations are in agreement with prior ^23^Na-MRI results(Driver et al., 2020; Gilles et al., 2017; Huhn et al., 2019; Meyer et al., 2019; Ridley et al., 2018).

Administration of TmDOTP^5-^ resulted in the emergence of multiple ^23^Na peaks (**Figure 3, green spectra**), particularly within the tumor. However the positive downfield shifts seen in healthy tissue suggest the paramagnetic effects of TmDOTP^5-^ reached the extracellular space even with limited extravasation. We found the most shifted peak ~2 ppm, sufficiently far from the other two peaks present. Therefore, this peak can be confidently attributed to only Na^+^_b_, and its integral 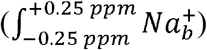 reflected the blood sodium concentration [Na^+^]_b_. Likewise 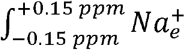 and 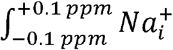 measured the [Na^+^]_e_ and [Na^+^]_i_, respectively. Tumor voxels [**Figure 3(a)**: voxels 4,5; **Figure 3(b)**: voxels 3,4] exhibited spectra where the three peaks were most notably present, with the most shifted peak occurring at ~2 ppm, the intermediate at ~0.5 ppm, and an unshifted peak at 0 ppm. Thus the chemical shifts of the Na^+^_b_, Na^+^_e_, and Na^+^_i_ peaks can be respectively placed at 2 ppm, 0.5 ppm and 0 ppm. Shifts of this nature were evident throughout the entire depth of the brain for both animals (**Figure S3**). Ventricular voxels [**Figure 3(a)**: voxels 1,6; **Figure 3(b)**: voxels 1,5] displayed only one Lorentzian peak shifted ~0.5 ppm. Healthy tissue voxels [**Figure 3(a)**: voxels 2,3; **Figure 3(b)**: voxels 2,6] also displayed one shifted peak ~0.5 ppm but with super-Lorentzian lineshape. This Na^+^_e_ shift in tissue coincides with the ventricular shift, as CSF and the extracellular space are physically in contact with unrestricted exchange of aqueous Na^+^. Given the shiftability *s*_[*paraCA^n−^*]_ is 2.77 ppm/mM measured *in vitro* (**Figure 1**), the tumor vasculature contained no more than 0.7 mM TmDOTP^5-^. Since the blood ^23^Na signal experiences the greatest shift, the (extracellular) tissue therefore encountered even less TmDOTP^5-^, in agreement with prior observations(Coman, Trubel, Rycyna, et al., 2009).

### *In Vivo* Depiction of Transmembrane and Transendothelial Na^+^ gradients

Integration of compartmentalized ^23^Na magnitude-corrected spectra (**Figures 2–3 and S2-S3**) generated spatial maps which showed relative [Na^+^] in each compartment from which the transmembrane (ΔNa^+^_mem_ = ∫Na^+^_e_ - ∫Na^+^_i_) and transendothelial (ΔNa^+^_end_ = ∫Na^+^_b_ - ∫Na^+^_e_) gradient maps could also be calculated, as shown in **Figure 4** for multiple axial slices bearing an RG2 tumor. This 3D high-resolution demonstration of the *in vivo* Na^+^ biodistribution divulges spatial heterogeneity, where the relative [Na^+^] of each compartment is a function of the compartment volume and the amount of Na^+^ in that compartment.

**Figure 4.**
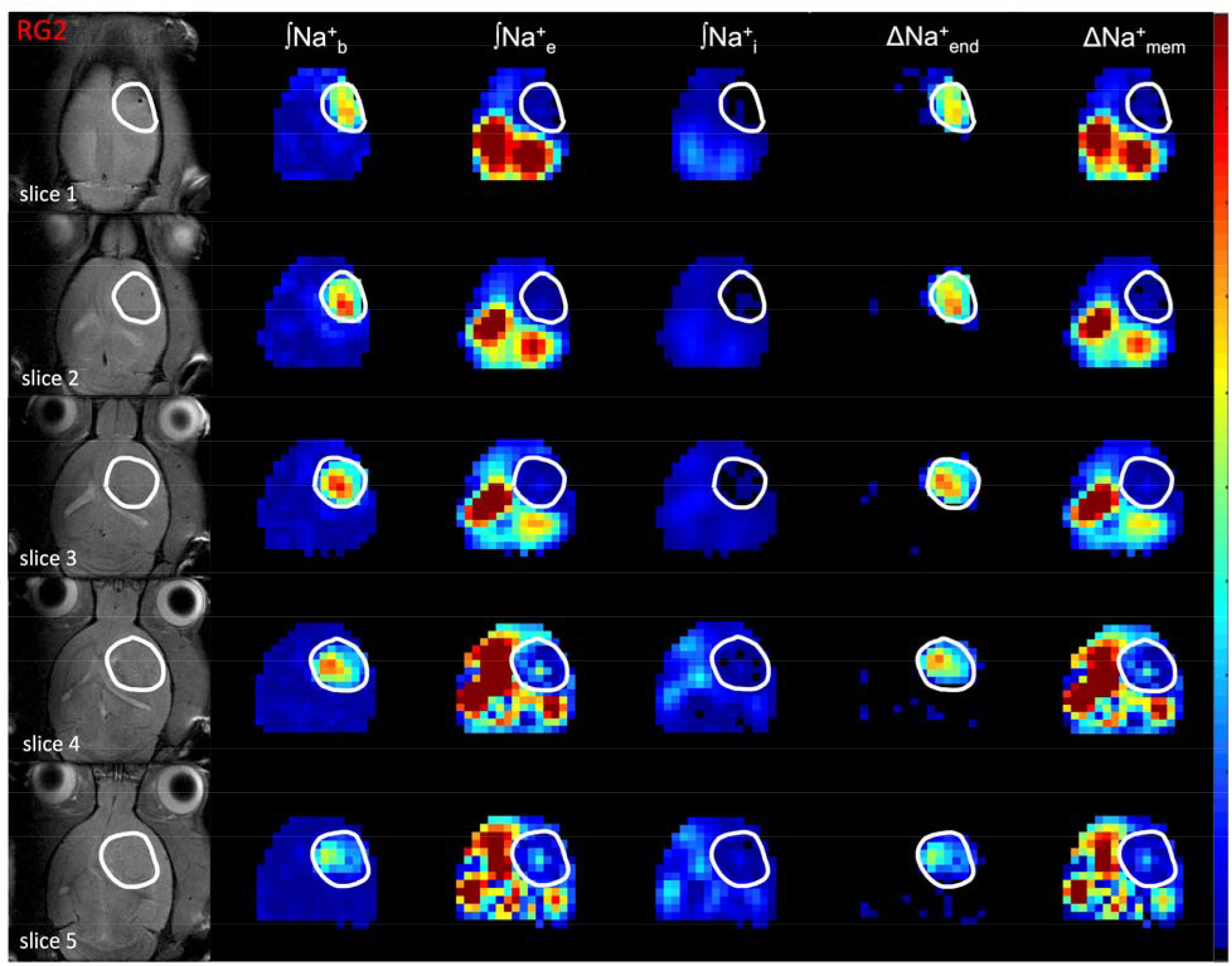
Spatial distributions of compartmentalized ^23^Na signals (Na^+^_b_, Na^+^_e_, Na^+^_i_) as well as transendothelial (ΔNa^+^_end_) and transmembrane (ΔNa^+^_mem_) gradients in an RG2 tumor. The left column shows the tumor location (white outline) on the anatomical ^1^H-MRI. Since the integral of each ^23^Na peak represents the [Na^+^], the respective three columns show the integral maps of Na^+^_b_, Na^+^_e_, and Na^+^_i_ from left to right (i.e., ∫Na^+^_b_, ∫Na^+^_e_, ∫Na^+^_i_). The last two columns on the right show ΔNa^+^_end_ = ∫Na^+^_b_-∫Na^+^_e_ and ΔNa^+^_mem_ = ∫Na%-∫Na^+^_i_. The ∫Na^+^_b_ map reveals low values in healthy tissue compared to tumor tissue, and within the tumor boundary a high degree of heterogeneity. The ∫Na^+^_e_ map reveals low values in tumor and normal tissues, but within the tumor boundary a small degree of heterogeneity is visible while ventricular voxels show very high values. The ∫Na^+^_i_ map reveals low values ubiquitously except some ventricular voxels. The ΔNa^+^_end_ map reveals dramatically high values within the tumor only. The ΔNa^+^_end_ was driven primarily by an increase of ∫Na^+^_b_ inside the tumor and which was more pronounced in superficial regions of the brain compared to deeper slices. The ΔNa^+^_mem_ map shows low values in tumor tissue compared to normal tissue, although ventricular voxels show very high values. The ΔNa^+^_mem_ is driven primarily by decreased ∫Na^+^_e_ and thus shows similar level of heterogeneity as the ∫Na^+^_e_ map. All maps use the same color scale and are relative. See **Figure S4** for an example for a U87 tumor.

There was markedly increased ∫Na^+^_b_ in the tumor, which was not observed elsewhere in normal brain. There was also high degree of heterogeneity within the tumor. The ∫Na^+^_e_ map revealed the largest values in the ventricles (CSF) and smaller values in the tumor with a slight extent of heterogeneity. Outside the tumor, the bulk peak occurred in the region of integration for Na^+^_e_. The ∫Na^+^_i_ map unsurprisingly showed values that were about one order of magnitude lower throughout the brain compared to the ∫Na^+^_b_ and ∫Na^+^_e_ maps, since [Na^+^]_i_ (~10 mM) is an order of magnitude smaller than [Na^+^]_b_ and [Na^+^]_e_ (~150 mM). Furthermore, the ∫Na^+^_i_ values were not significantly different between the tumor and healthy tissue.

The ΔNa^+^_mem_ values in the tumor were significantly lower compared to the healthy tissue (*p* < 0.05) and the map displayed a similar level of heterogeneity as the ∫Na^+^_e_ map, suggesting that ΔNa^+^_mem_ is driven primarily by the decrease in Na^+^_e_. Ventricular voxels still showed high values in ΔNa^+^_mem_, indicating the large magnitude of Na^+^_e_ in CSF. Likewise, the significant elevation of ΔNa^+^_end_ in the tumor was driven primarily by the Na^+^_b_ increase, and ΔNa^+^_end_ values were significantly larger in the tumor compared to healthy tissue (*p* < 0.05). This was more pronounced in superficial regions of the brain. For both gradients, statistical significance was achieved even after excluding ventricle values. These patterns could also be visualized by looking at slice projections of the compartmental and gradient values for the same RG2-bearing animal along a constant coronal position (**Figure 5**). For the RG2 tumor, the tumoral increases in Na^+^_b_ and ΔNa^+^_end_ were highest superficially (slices 1-4). Conversely, peritumoral values of Na^+^_e_ and ΔNa^+^_mem_ increased with depth up to a point in the middle of the brain (slices 3-4) before diminishing. Intratumoral Na^+^_e_, however, did not vary significantly with depth. Na^+^_i_ also decreased inside the tumor but not significantly. The ΔNa^+^_mem_ and ΔNa^+^_end_ respectively behaved similarly to Na^+^_b_ and Na^+^_e_ since they were the primary drivers of those gradients. Similar observations were made for U87 tumors (**Figures S4-S5**) regarding Na^+^ in each compartment and the corresponding gradients.

**Figure 5.**
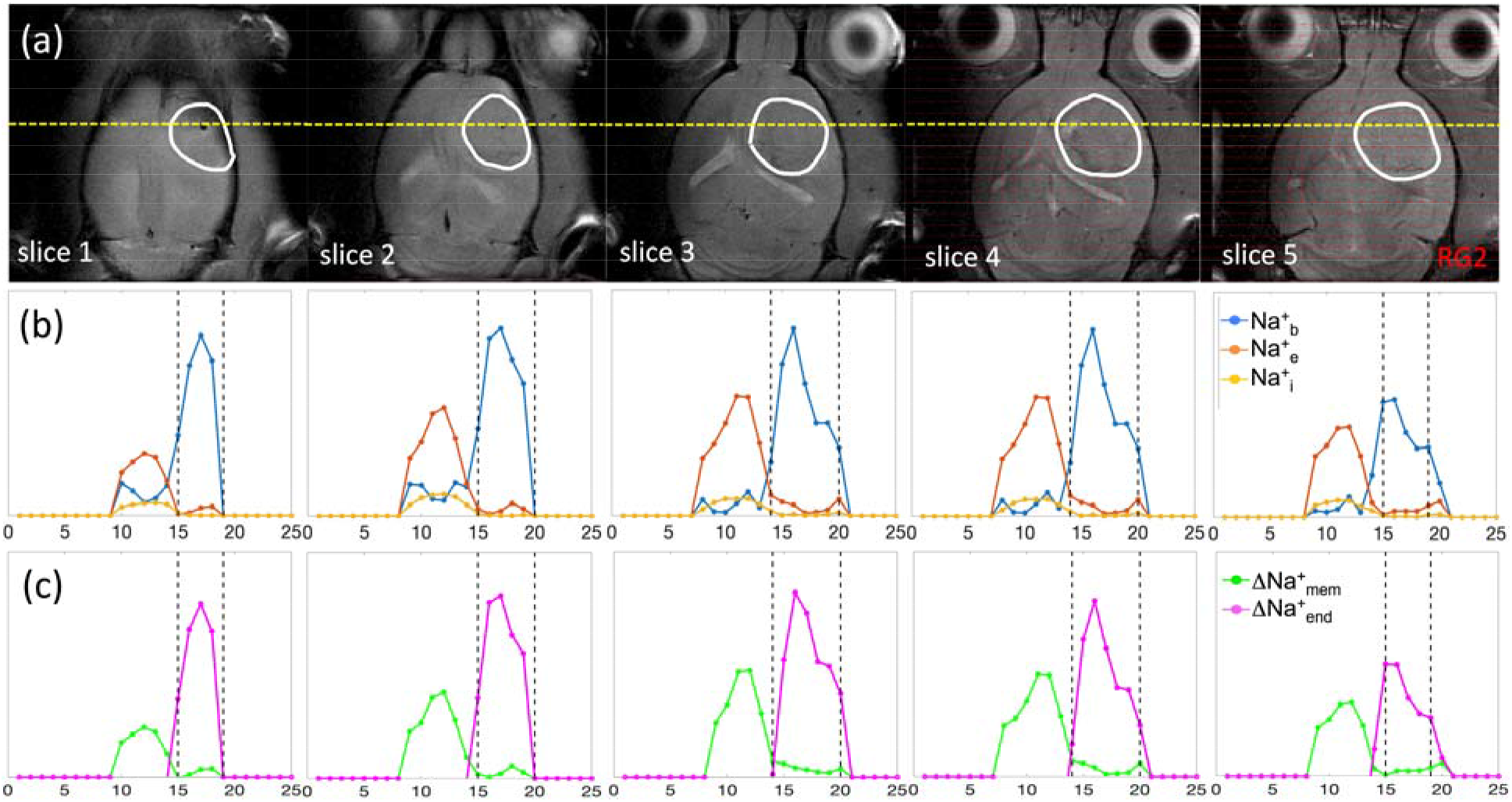
Coronal projections of compartmentalized ^23^Na signals (Na^+^_b_, Na^+^_e_, Na^+^_i_) as well as transendothelial (ΔNa^+^_end_) and transmembrane (ΔNa^+^_mem_) gradients in an RG2 tumor. (a) Axial ^1^H-MRI indicating the tumor (white outline) across slices (same as **Figure 4**), where the yellow line indicates the position for a coronal projection. (b) Spatially varying ^23^Na signals for Na^+^_b_, Na^+^_e_, and Na^+^_i_ are shown with blue, orange, and yellow lines, respectively, where the vertical black lines indicate the tumor boundary. The Na^+^_b_ signal (blue) is clearly elevated in the tumor, and most elevated in slices 1-4 (or superficially). Behavior of Na^+^_b_ signal (blue) is inversely related to Na^+^_e_ signal (orange), which is high outside the tumor and weaker inside the tumor. While intratumoral Na^+^_b_ signal (blue) is high in slices 1-4, the peritumoral Na^+^_e_ signal (orange) is highest in slices 3-4. Comparatively, the Na^+^_i_ signal (yellow) does not vary significantly across slices, but slightly lower inside the tumor than outside the tumor. (c) Behaviors of ΔNa^+^_mem_ (green) and ΔNa^+^_end_ (magenta) signals closely mimic patterns of Na^+^_e_ and Na^+^_b_ signals, respectively, indicating that each of those Na^+^ compartments is the primary driver of the respective Na^+^ gradient. See **Figure S5** for a similar example for a U87 tumor.

Throughout the entire cohort of animals (**Figure 6(a,b)**), the mean ∫Na^+^_b_ values were larger and ∫Na^+^_e_ values were lower in the tumor compared to normal tissue. These trends were significant in RG2 (*p* < 0.005) and U87 (*p* < 0.05) tumors while there was no significant difference in ∫Na^+^_i_ for all three tumors (**Figure 6(a)**). Identical trends were also observed in ΔNa^+^_end_ and ΔNa^+^_mem_ values, and significantly so in RG2 (*p* < 0.005) and U87 (*p* < 0.05) tumors. Moreover, ΔNa^+^_end_ was significantly stronger in RG2 and U87 tumors compared to U251 (*p* < 0.05) (**Figure 6(b)**).

**Figure 6.**
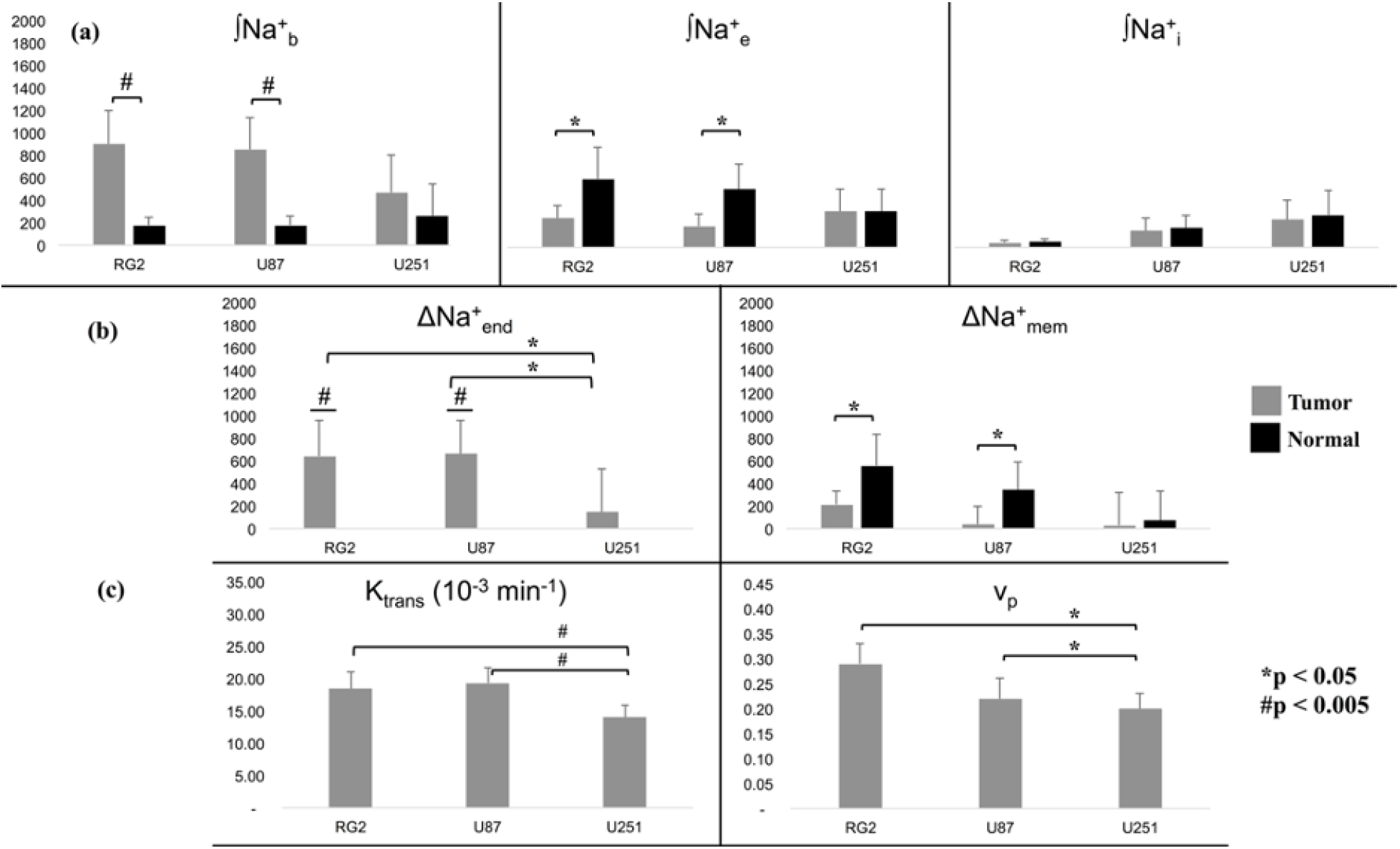
Statistical comparisons between intracellular, extracellular, and vascular compartments across RG2, U87, and U251 tumors with ^23^Na-MRSI and ^1^H-DCE-MRI. (a) Relation between ∫Na^+^_b_, ∫Na^+^_e_ and ∫Na^+^_i_ across tumor and healthy tissues. For the RG2 and U87 tumors, the ∫Na^+^_b_ values were significantly higher than normal tissue (*p* < 0.005, #). Also for these tumors, the ∫Na^+^_e_ values were significantly lower than normal tissue (*p* < 0.05, *). The mean values for the U251 tumor roughly followed the same trend but were not significant. Furthermore, there was no significant difference between ∫Na^+^_i_ values in tumor and normal tissues for any of the three tumor types. (b) Relations between tumor and normal tissues for ΔNa^+^_en_d and ΔNa^+^_mem_ for the three tumor types. Tumor ΔNa^+^_end_ values were significantly larger than normal values (*p* < 0.005, #), which were non-positive (data not shown). Moreover, ΔNa^+^_end_ in RG2 and U87 tumors was significantly greater than in the U251 tumor (*p* < 0.05, *), indicative of vascular differences between the tumor types. ΔNa^+^_mem_ values were, on average, weaker in tumor compared to normal tissue, but significant only in RG2 and U87 tumors (*p* < 0.05, *). Based on **Figures 5 and S5**, it is clear that the relation between ΔNa^+^_end_ and ΔNa^+^_mem_ is negative. (c) ^1^H-DCE-MRI data for *K^trans^* and *v_p_* values, which are known to reveal information regarding vascular structure and function. *K^trans^* follows the same patterns as ∫Na^+^_b_ and ΔNa^+^_end_ across tumor types. *K^trans^* (*p* < 0.005, #) and *v_p_* (*p* < 0.05, *) were both significantly larger in RG2 and U87 tumors, compared to U251. See **Figure S6** for the *F_p_* and *v_e_* ^1^H-DCE-MRI parameters for each tumor type. See **Figure S7** for exemplary maps of ^1^H-DCE-MRI parameters for individual animals from each tumor type.

Since a strengthened ΔNa^+^_end_ is indicative of impaired vascular integrity, we employed ^1^H-DCE-MRI to reliably image vascular dynamics and function(Sourbron & Buckley, 2013). Of the four parameters which can be obtained by fitting ^1^H-DCE-MRI data from a 2XCM, the volume transfer constant (*K^trans^*) and plasma volume fraction (*v_p_*), as shown in **Figure 6(c)**, both followed the trends of ΔNa^+^_end_ across tumor types: in RG2 and U87 tumors compared to U251, there was a significant difference (*K^trans^*: *p* < 0.005 and *v_p_*: *p* < 0.05) (for plasma flow rate (*F_p_*) and extracellular volume fraction (*v_e_*) see **Figure S6**). Although significance was marginal for *F_p_*, the mean values followed suit (**Figure S6**). The ^1^H-DCE-MRI data displayed a region of simultaneous low *F_p_* and larger *v_e_* within an exemplary slice of a U251 tumor, indicative of a necrotic core, which RG2 and U87 animals lacked (**Figure S7**). Additionally, *v_e_* on average was smaller than *v_p_*, indicating a high degree of vascularity/angiogenesis in tumors. These findings further substantiate the ∫Na^+^_b_ and ΔNa^+^_end_ results derived from the ^23^Na-MRSI studies.

**Figure 7** shows compartmental and gradient maps across all tumor cell lines (RG2, U87, U251). The trends seen previously pervaded all animals, but to varying degrees based on the tumor type. The Na^+^_b_ elevation, and concomitant ΔNa^+^_end_ strengthening, were most pronounced for the RG2 tumor, followed by U87 and then U251. The heterogeneity of ∫Na^+^_b_ and ΔNa^+^_end_ also followed the same order, as did the decrease in ∫Na^+^_e_ and weakening of ΔNa^+^_mem_. In all tumors, Na^+^_b_ and Na^+^_e_ patterns respectively drove the behaviors of ΔNa^+^_end_ and ΔNa^+^_mem_.

**Figure 7.**
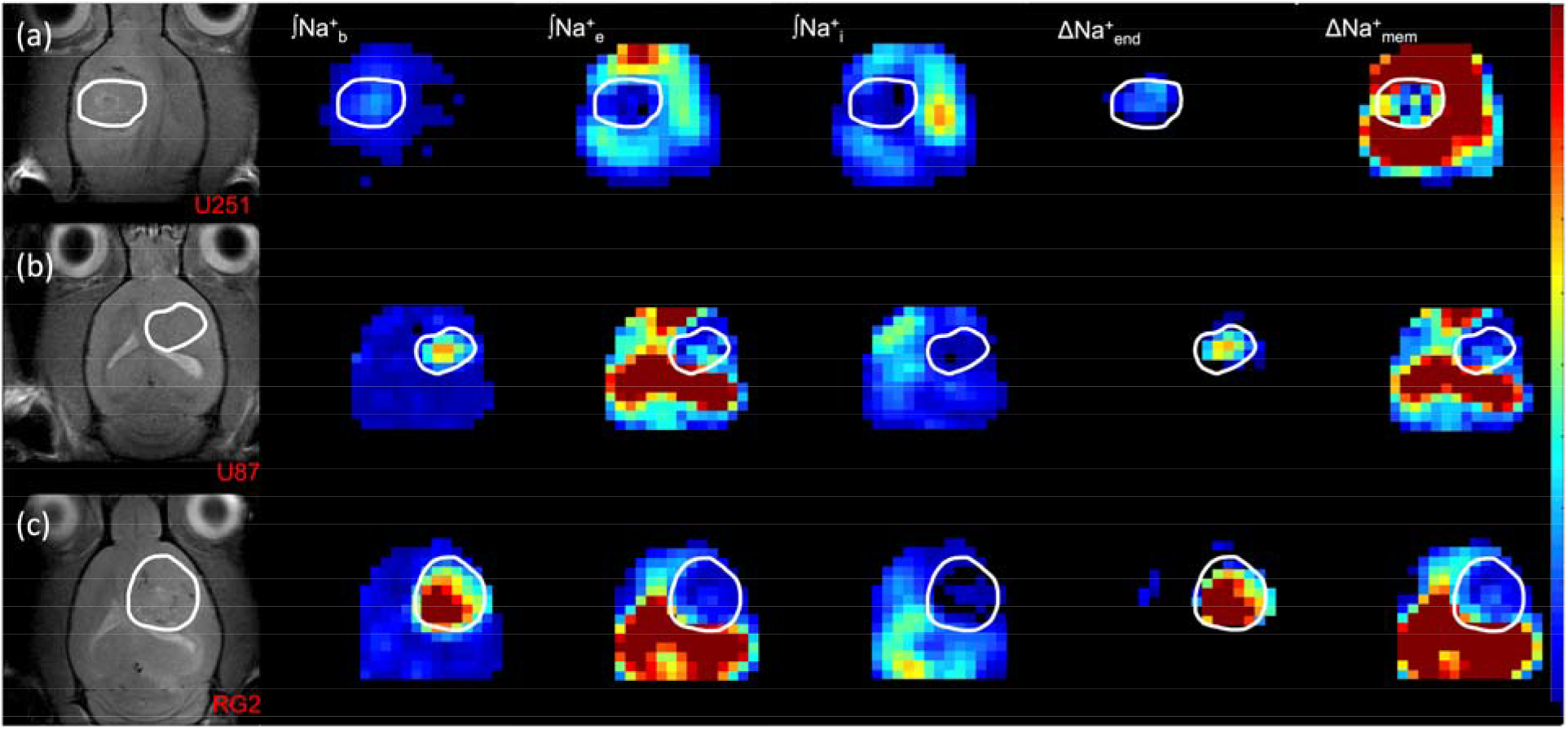
Representative maps of compartmentalized ^23^Na signals (Na^+^_b_, Na^+^_e_, Na^+^_i_) as well as transendothelial (ΔNa^+^_end_) and transmembrane (ΔNa^+^_mem_) gradients in U251, U87, and RG2 tumors. The left column shows the tumor location (white outline) on the anatomical ^1^H-MRI for animals bearing (a) U251, (b) U87 and (c) RG2 tumors. The respective three columns show ∫Na^+^_b_, ∫Na^+^_e_, and ∫Na^+^_i_ maps. The last two columns on the right show the ΔNa^+^_end_ and ΔNa^+^_mem_ maps. In all tumors the ∫Na^+^_b_ and ∫Na^+^_e_ are high and low, respectively, and thus are the main drivers for a high ΔNa^+^_end_ and a low ΔNa^+^_mem_.

## DISCUSSION

### Study Highlights

*In vitro* ^23^Na shifts were most dependent on [TmDOTP^5-^] given its high shiftability (*s*_[paraCA^*n*−^]_ =2.77 ppm/mM), whereas shiftability due to pH and temperature effects were negligible within physiological ranges (*s_pH_*=0.25 ppm/pH unit; *S_T_*=0.03 ppm/°C). The maximum pH difference between glioma and brain tissue is ~0.4 pH units(Coman et al., 2016; Huang et al., 2016; Maritim et al., 2017; Rao et al., 2017) whereas temperature differences of ~0.5 °C are extremely unusual in the brain(Coman et al., 2013; Coman et al., 2015; Walsh et al., 2020). Under these extreme conditions, the respective ^23^Na shift variations caused by pH and temperature would be 0.1 ppm and 0.015 ppm. Meanwhile, TmDOTP^5-^ can reach *in vivo* concentrations close to 1-2 mM in blood and interstitial spaces(Coman, Trubel, & Hyder, 2009; Coman, Trubel, Rycyna, et al., 2009; Trübel et al., 2003) which would cause ^23^Na shifts of 2.8-5.5 ppm. Given observed ^23^Na line widths *in vivo* on the order of ~0.4 ppm, TmDOTP^5-^ concentration effects dominate the shifting effect (96-98%). Therefore, ^23^Na shiftability can be considered a univariate function of [TmDOTP^5-^] *in vivo*.

This observation enabled attributing individual ^23^Na peaks to specific *in vivo* pools for blood, extracellular and intracellular spaces arising from compartmental differences in [TmDOTP^5-^] upon intravenous administration. The shifts in tumor tissue were more conspicuous compared to peritumoral tissue but where [TmDOTP^5-^] was lower (~1 mM) than in blood (~2 mM). Additionally, the blood and extracellular peaks were separated by ~1.5 ppm, much larger than their line widths (~0.4 ppm), indicating minimal cross-compartmental contributions.

Integrating the separated ^23^Na peaks enabled spatial mapping of Na^+^ compartments and gradients for the first time *in vivo*. In the tumor, compared to normal tissue, the transendothelial Na^+^ gradient was stronger and the transmembrane Na^+^ gradient was weaker due to elevated blood and decreased extracellular ^23^Na signals. The enhanced ^23^Na blood signals in tumors complied with dynamic ^1^H-DCE-MRI scans based on gadolinium (Gd^3+^) uptake, which revealed a higher degree of vascularity in RG2 and U87 tumors. Extracellular Na^+^ signal in the ventricles was also very high due to the dominance of CSF. However, ventricular ^23^Na peaks were Lorentzian whereas tissue ^23^Na peaks appeared super-Lorentzian, since CSF contains only aqueous Na^+^ and thus a single *T_2_* component, whereas semi-solid Na^+^ binding in tissue resulted in multiple *T_2_* components(Sinclair et al., 2010).

### Comparison with Previous Work

The present *in vitro* data improve upon earlier attempts at quantifying ^23^Na shiftability using paraCA^*n*-^ versus many parameters like pH, temperature and other cations(Puckeridge et al., 2012). However, the findings focused more on characterizing the dependence on each parameter (linear, sigmoidal, etc.) rather than considering relevant *in vivo* conditions. Additionally, the model was not employed in the context of the brain/other tissues. Our ^23^Na shiftability model does not require assessing the effects of cationic competition for attraction to TmDOTP^5-^(Ren & Sherry, 1996) because other cations are not present in the blood and/or extracellular spaces in concentrations comparable to Na^+^(Cheng et al., 2013; Janle & Sojka, 2000; Romani, 2011).

Prior *in vivo* ^23^Na spectroscopy studies utilizing TmDOTP^5-^ in the brain failed to elucidate spatial information by only focusing on global data acquisition and/or localized voxels(Bansal et al., 1992; Winter et al., 1998). The findings reported two broadened peaks, an unshifted intracellular peak and a shifted extracellular peak. Based on two peaks over limited spatial regions, these studies could not comment specifically on the transmembrane gradient. Furthermore, the blood ^23^Na signal could not be separated fully so statements about the transendothelial gradient could not be made. The shifting capability of TmDOTP^5-^ for separating ^23^Na resonances in tumor tissue was demonstrated *in situ*, but still at a global level and without mention of Na^+^_b_ specifically(Winter & Bansal, 2001; Winter et al., 2001).

Recently, ^23^Na-MRI methods have been preferred clinically(Madelin, Lee, et al., 2014). Such relaxometric modalities exploit differences in diffusion and relaxation behavior between Na^+^ ions inside/outside the cell, because intracellular ions are generally considered less mobile due to binding. Due to the spin-3/2 of ^23^Na, this binding amplifies the relative contribution of nuclear satellite transitions and permits the use of MQF techniques to isolate signals from individual *in vivo* compartments. However, these ^23^Na-MRI methods need specificity for intracellular Na^+^ because they fail to completely suppress ^23^Na signals from the blood and extracellular compartments(Madelin, Lee, et al., 2014). Moreover, γ_Na_ is about one-quarter of γ_H_, which impairs sensitivity. These methods also employ large, fast-switching gradients.

Our method obviates these practical limitations and still provides relevant physiological information. Overall, the ^23^Na-MRSI results agree with prior findings that a depolarized *V_m_* (i.e., weakened transmembrane gradient) is responsible for tumor proliferation(Yang & Brackenbury, 2013). Given that both the cell membrane and BBB help to maintain the ionic level of the extracellular fluid(Bean, 2007; Ennis et al., 1996), our results also show that the transendothelial gradient is significantly enhanced in the same tumors that show enhanced permeability (i.e., RG2 and U87). Together these suggest that the current ^23^Na-MRSI scheme can be used to study the perturbed sodium homeostasis *in vivo* within the tumor neurovascular unit.

### Technical Limitations

This technique, while a crucial first step toward non-invasively mapping the spatial distribution of Na^+^ *in vivo*, cannot absolutely quantify [Na^+^]. This limitation can usually be circumvented by including a quantifiable standard, but ^23^Na-NMR has no species that can be used as a standard. However, using the strong CSF signal *in vivo* remains a possibility for future explorations. Setups involving Na^+^ phantoms with relatively large [TmDOTP^5-^] within the FOV alongside the body region being imaged could be used, but they hinder the shim around the subject’s body part and are difficult to cover with surface coils. Additionally, broad point-spread functions make quantifications in external phantom standards challenging, though they are perhaps the best option presently(Thulborn et al., 2019).

Contrast agents with lanthanide(III) ions (Ln^3+^) are popular in molecular imaging with ^1^H-MRI(Hyder & Manjura Hoque, 2017; Sherry & Woods, 2008), but clinically the preference is probes with Gd^3+^ conjugated to linear or cyclical chelates(Herborn et al., 2007; Kubicek & Toth, 2009). The most biocompatible Gd^3+^ chelates are based on 1,4,7,10-tetraazacyclododecane-1,4,7,10-tetraacetate (DOTA^4-^) because they are both kinetically and thermodynamically stable(Sherry et al., 2009). A LnDOTA^-^ carries a −1 charge. But if a phosphonate group is attached to each of the pendant arms in DOTA^4-^, then DOTP^8-^ is formed and complexation with Ln^3+^ permits a −5 charge (e.g. TmDOTP^5-^). The majority of paraCA^*n*-^ that will work for the type of ^23^Na-MRSI experiments described here are based on Ln^3+^ complexes, because these give rise to large hyperfine shifts with minimal ^1^H relaxation enhancement(Huang et al., 2015). But there is growing attention to complexes with similar paramagnetic properties from transition(II) metal ions (Tn^2+^), such as Fe^2+^, Ni^2+^, or Co^2+^(Tsitovich & Morrow, 2012). Tn^2+^-based paraCA^*n*-^ have the potential for clinical use because of superior biocompatibility. Some Tn^2+^ complexes designed could carry a −5 charge also, but studies need to explore the safest and most effective paraCA^*n*-^ for ^23^Na-MRSI experiments.

Another limitation is the infusion of a small amount of Na^+^ with the paraCA^*n*-^ itself. TmDOTP^5-^ exists commercially in the form Na_5_TmDOTP, so a small amount of Na^+^ is being added. Since [TmDOTP^5-^] does not exceed 2 mM in the brain vasculature, there is at most ~1.3% increase of the endogenous [Na^+^] in blood/extracellular spaces. In regions with high [TmDOTP^5-^]/low [Na^+^], like the necrotic core of tumors, the infused Na^+^ may represent a larger percentage. However, necrotic cores can be identified with *T_2_*-weighted ^1^H-MRI scans. Since extracellular Na^+^ is shifted less than blood, it is doubtful that enough Na_5_TmDOTP extravasation occurs to significantly alter the relative Na^+^ levels between compartments and impact the conclusions drawn from this study. Future studies with localized opening of the BBB at higher magnetic fields can help with these uncertainties.

### Implications of Current Findings

Our results enabled comparisons of Na^+^ physiology and distributions among RG2, U87, and U251 gliomas. Both U87 and U251 are human-derived cell lines, whereas RG2 is derived from rat glioma(Aas et al., 1995; Jiang et al., 2018). Experimentally the U251 tumor is most heterogeneous, since U251 cells grow erratically and anisotropically compared with RG2 and U87 cells. The U251 tumor is more invasive and infiltrative than U87(Candolfi et al., 2007). U251 cells display greater necrosis, expression of hypoxia-inducible factor 1-alpha (HIF1α) and of Ki67, indicating higher rates of proliferation(Radaelli et al., 2009). Cells also test positive for glial fibrillary acidic protein (GFAP) and vimentin, and exhibit neovascularization and angiogenesis. U87 cells are also positive for vimentin and exhibit significant angiogenesis but do not develop necrosis. Neither U251 nor U87 exhibits endothelial proliferation, a common hallmark of human-derived GBM lines(Candolfi et al., 2007). The RG2 tumor exhibits invasiveness and induces BBB disruption, producing edema surrounding the tumor where pericytes help promote angiogenesis to increase permeability of the tumor vasculature(Hosono et al., 2017). These data concur with our findings. We observed that the negative correlation between the transmembrane and transendothelial gradients were strong in the RG2 and U87 lines but weak for U251. The increase of the transendothelial gradient nearly matched the decrease of the transmembrane gradient in U87 tumors, and exceeded in RG2, which matched behavior regarding BBB permeability. Higher density of blood vessels or higher blood volume would explain higher ^23^Na signal but not necessarily higher Na^+^ concentration in the blood. Although the blood vessels are leaky to Gd^3+^, the elevated transendothelial gradient suggests that the BBB is impermeable to Na^+^, which is well known(Ennis et al., 1996).

Alkylating chemotherapy agents attach an alkyl group to DNA of cancer cells to keep them from replicating. For example, temozolomide (TMZ) achieves cytotoxicity by methylating the O^6^ position of guanine. O^6^-methylguanine-DNA-methyltransferase (MGMT) is a DNA repair enzyme, which ordinarily repairs the naturally occurring DNA lesion O^6^-methylguanine back to guanine and prevents mistakes during DNA replication and transcription. Unfortunately, MGMT can also protect tumor cells by the same process and neutralize the cytotoxic effects of agents like TMZ. If the *MGMT* gene is silenced by methylation in tumor cells (i.e. MGMT-negative or MGMT-methylated), its DNA repair activity is diminished and the tumor’s sensitivity to chemotherapy is amplified. This suggests that MGMT-positive tumor cells become resistant to chemotherapy, and therefore would possess a depolarized *V_m_* due to its proliferative state.

A recent study demonstrated higher MGMT mRNA expression for RG2 compared to U87(Lavon et al., 2007). Another study showed that the 50% inhibition concentration (IC^50^) of TMZ for U87 and U251 cells are comparable(Qiu et al., 2014). Together, these suggest that RG2 is most resistant to chemotherapy presumably due to its augmented proliferative/replicative state, and hence a depolarized *V_m_*. These observations partially agree with our results, where RG2 and U87 tumors maintain a depolarized *V_m_* for their proliferative/replicative state to persist.

## CONCLUSION

This study is the first to non-invasively image the transformed transmembrane and transendothelial gradients of gliomas using TmDOTP^5-^ for 3D ^23^Na-MRSI at high spatial resolution. The *in vivo* data consistently revealed ΔNa^+^_mem_ weakening and ΔNa^+^_end_ strengthening within tumors compared to normal tissue, which agree with prior findings(Yang & Brackenbury, 2013) and suggest a redistribution of tumoral Na^+^_e_ to the blood compartment. There is good evidence to propose that these measurements could potentially probe stages of the cell cycle, and perhaps, angiogenic behavior. *In vivo* testing of novel chemotherapy and anti-angiogenic drugs for GBM models even at a preclinical level would be significant. However, this method could potentially be translated into patients by synthesizing transition metal-based paraCA^*n*-^ such that suitable therapies can be targeted based on MGMT screening in GBM patients.

## MATERIALS AND METHODS

### *In vitro* characterization

*In vitro* experiments were performed using a 2-compartment coaxial cylindrical 7-inch NMR tube setup from WilmadLabGlass (Vineland, NJ, USA). One compartment contained 150 mM NaCl and the other contained the same but with varying amounts of TmDOTP^5-^ (1–10 mM) and 10% v/v ^2^H_2_O to lock the spectrometer frequency using the ^2^H_2_O signal (**Figure S1**). NaCl and H_2_O were purchased from Sigma-Aldrich (St. Louis, MO, USA), and TmDOTP^5-^ was purchased as the sodium salt Na_5_TmDOTP from Macrocyclics (Plano, TX, USA). The 5-mm opening of the NMR tube permitted an insert (the inner compartment) whose 50-mm-long tip had inner and outer diameters of 1.258 and 2.020 mm, respectively. The outer-to-inner volume ratio between the two compartments was 8.6. The geometry of the setup allowed 645 μL total in the outer compartment to fill around the tip. Each solution was pH-adjusted using HCl or NH_4_OH to give 5 different pH values.

^23^Na NMR spectra were collected on a Bruker Avance III HD 500 MHz vertical-bore spectrometer (Bruker, Billerica, MA, USA) interfaced with Bruker TopSpin v2.1 software. A single ^23^Na square pulse (50 μs) was used to globally excite the volume of interest (repetition time *T_R_*=275 ms) collecting 2048 free induction decay (FID) points in the time domain with an acquisition time *t*_aq_=38.9 ms, averaged 4096 times. Each set of scans was repeated at a series of temperatures: 27, 30, 34, 37, and 40 °C. Spectra were analyzed using 10-Hz line broadening and manual zeroth- and first-order phasing.

### *In vivo* studies

The *in vivo* protocol was approved by the Institutional Animal Care & Use Committee of Yale University. Rats (athymic/nude and Fischer 344) were purchased through Yale University vendors. U251, U87 and RG2 GBM cell lines were purchased from American Type Culture Collections (Manassas, VA, USA). The U251, U87, and RG2 cells were cultured and grown in a 5% CO_2_ incubator at 37 °C in either low-glucose (U251 cells) or high-glucose (U87 and RG2 cells) Dulbecco’s Modified Eagle’s Medium (DMEM) (Thermo Fisher Scientific, Waltham, MA, USA) with 10% fetal bovine serum (FBS) and 1 % penicillin-streptomycin. Cells for tumor inoculation were harvested upon reaching at least 80% confluence and were prepared in FBS-free DMEM. Athymic/nude rats were injected intracranially with 2–5×10^6^ tumor cells either from the U251 (*n*=6) or the U87 (*n*=8) cell line (5-μL aliquot) while placed in a stereotactic holder on a heating pad. Fischer 344 rats were injected with 1.25×10^3^ RG2 cells (*n*=8). During the procedure, animals were anesthetized via isoflurane (Isothesia™) inhalation (3–4%), purchased from Covetrus (Portland, ME, USA). Injections were performed using a 10-μL Hamilton syringe with a 26G needle into the right striatum for majority of the experiments, 3 mm to the right of the bregma and 3 mm below the dura. The cells were injected steadily at 1 μL/min over 5 minutes and the needle was left in place for an additional 5 minutes post-injection. The syringe was then gradually removed to preclude any backflow of cells. The hole in the skull was sealed with bone wax, and the incision site was sutured after removal of the syringe. Animals were given bupivacaine (2 mg/kg at incision site) and carprofen (5 mg/kg, subcutaneously) during the tumor inoculation to relieve pain. Carprofen was subsequently given once per day for two days post-inoculation.

Rats were weighed daily and kept on a standard diet of rat chow and water. Tumor growth was monitored regularly using ^1^H-MRI. When the tumor had reached a minimum mean diameter of 3 mm, each animal was imaged using ^1^H-MRI and ^23^Na-MRSI. This generally occurred around 21 days post-injection. An infusion line was first established through cannulation of the tail vein as a means to administer fluids and the paraCA^*n*-^. During the cannulation procedure, the rat was placed on a heating pad to maintain physiological body temperature. A 30G needle, fitted onto a PE-10 line, was inserted into the tail vein while the animal was under anesthesia. The animal was then given Puralube Vet Ointment (Dechra, Overland Park, KS, USA) over the eyes and then situated in a prone position underneath a ^23^Na/^1^H quad surface coil before being placed in the magnet. The 2.5-cm ^23^Na coil was placed directly on top of the head, and the two 5-cm ^1^H coils flanked the head on the left and right sides. Breathing rate was measured by placement of a respiration pad under the torso, and temperature was monitored through a rectal fiber-optic probe thermometer.

Imaging was conducted on a 9.4T horizontal-bore Bruker Avance system, interfaced with Bruker ParaVision software running on CentOS. Positioning and power optimizations for ^1^H signals were performed using Bruker-defined gradient-echo (GE) and fast spin-echo (FSE) sequences. Shimming was done on the ^1^H coils using an ellipsoid voxel to bring the H_2_O line width to less than 30 Hz. Pre-contrast ^1^H anatomical MRI was first performed using a spin-echo sequence with 9 axial slices (field of view (FOV): 25×9×25 mm^3^, 128×128 in-plane resolution) over 10 echo times *T_E_* (10-100 ms) with *T_R_*=4 s. The multiple echo times enabled voxel-wise calculations of ^1^H *T*_2_ values. ^23^Na power optimizations were then performed using a global 2-ms Shinnar-Le Roux (SLR) pulse over an 8-kHz bandwidth (v_0_^Na^==105.9 MHz at 9.4 T). The optimal 90°-pulse power was achieved using no more than 8 W.

^23^Na-MRSI was performed using the same SLR pulse over a 25×19×25 mm^3^ FOV using a nominal voxel size of 1.0×1.0×1.0 mm^3^ with *T_R_*=300 ms, and phase encoding (gradient duration of 1 ms) done in all three spatial dimensions to avoid chemical shift artifacts caused by slice-selective radiofrequency pulses. This was done with reduced spherical encoding and a *k*-space radius factor of 0.55. A preliminary ^23^Na-MRSI scan was run before administering paraCA^*n*-^. The *in vivo* ^1^H-MRI delineated the tumor and brain boundary and permitted co-registration with ^23^Na-MRSI data, both before and after infusion of paraCA^*n*-^, enabling anatomical localization of ^23^Na-MRSI spectra at the voxel level.

The animals were then given 1 μL/g body weight (BW) probenecid using a syringe pump (Harvard Apparatus, Holliston, MA, USA) for 10 minutes followed by a 20-minute waiting period. Then Na_5_TmDOTP (1 μmol/g BW) was co-infused with probenecid (same dose) at a rate of 15 μL/min. Post-contrast ^23^Na-MRSI was performed 30 minutes after the start of infusion and repeated subsequently thereafter during the infusion. The imaging session was concluded with post-contrast ^1^H-MRI under identical conditions.

^1^H-MRI and ^23^Na-MRSI results were processed and analyzed using home-written code in MATLAB (MathWorks, Natick, MA, USA). Voxel-wise *T*_2_ values for ^1^H were calculated by fitting MRI voxel intensities versus the series of *T_E_* values to a monoexponential curve *e^-TE/T2^*. Pre-contrast and post-contrast ^1^H *T*_2_ values were used to qualitatively ascertain the success of paraCA^*n*-^ infusion. 3D ^23^Na-MRSI data were reconstructed using Fourier transformation in all spatial and temporal dimensions after 10-Hz line-broadening. Integration of individual ^23^Na peaks was performed over the following ranges: (i) 2±0.25 ppm for Na^+^_b_, (ii) 0.5±0.15 ppm for Na^+^_e_, and (iii) 0±0.1 ppm for Na^+^_i_. Due to line broadening induced by TmDOTP^5-^, the integration range was different for each compartment to capture the complete integral. The ΔNa^+^_mem_ values were calculated by subtracting ∫Na^+^_i_ from ∫Na^+^_e_, and ΔNa^+^_end_ by subtracting ∫Na^+^_e_ from ∫Na^+^_b_.

### ^1^H-DCE-MRI Studies

To measure vascular parameters [*K^trans^* (volume transfer coefficient, min^-1^), *F_p_* (plasma flow rate, min^-1^), *v_e_* (extracellular volume fraction, unitless), *v_p_* (plasma volume fraction, unitless)] from a two-compartment exchange model (2XCM), ^1^H-DCE-MRI was performed on a subset of RG2 (9.4T), U87 (11.7T) and U251 (11.7T) tumors. ^1^H-DCE-MRI data used a ^1^H volume-transmit (8-cm)/surface-receive (3.5-cm) coil.

Baseline images for *T_1_* mapping were acquired using a rapid acquisition with relaxation enhancement (RARE) sequence with six *T_R_* values (0.4, 0.7, 1, 2, 4, 8 s). Seven 1-mm slices covering the extent of the tumor were chosen and images were acquired with a 25×25 mm FOV, 128×128 matrix and *T_E_* of 10 ms. ^1^H-DCE-MRI acquisition consisted of a dynamic dual-echo spoiled GE sequence with a temporal resolution of 5 s. Images were acquired with *T_R_*=39.1 ms, *T_E_*=2.5/5 ms, flip angle=15°, and one average. Three central slices of the tumor were chosen with identical positioning, FOV (25×25 mm), and matrix (128×128) to be co-registered to the *T_1_* data.

The sequence was repeated every 5 s over a 22 min period with 0.25 μmol/g gadobutrol (Bayer, AG), a gadolinium (Gd^3+^)-containing contrast agent, injected 2 min after the start of the sequence and then flushed with 100 μL heparinized saline. The multi-*T_R_ T_1_* sequence was then repeated at the end of the ^1^H-DCE-MRI acquisition to serve as a post-Gd^3+^ *T_1_* mapping which was used to delineate tumor boundaries. Quantitative *T_1_* maps were generated by fitting voxel-level data to a monoexponential function in MATLAB.

Measurements from *T_1_*-weighted images before Gd^3+^ injection were used to transform time-intensity curves into time-concentration curves after the bolus injection. The region of interest (ROI) was placed inside the tumor area, including the rim, as determined by the region of contrast enhancement/uptake. All analysis, including masking the ROI, was performed in MATLAB using the same home-written code. The arterial input function (AIF) was measured by collecting arterial blood samples at discrete time points post-injection. The raw AIF was fit to a bi-exponential curve with a linear upslope during injection of Gd^3+^. Plasma [Gd^3+^] was derived from the blood [Gd^3+^] using a hematocrit of 0.45. The time resolution and duration interval used downstream in the analysis pipeline were adjusted manually.

The 2XCM parameters were estimated by fitting each voxel using Levenberg-Marquardt regression. Because *K^trans^* fitting often converged on local minima instead of the desired global minimum, multiple starting values were used, ultimately choosing the one with the smallest residual. Other variables were less sensitive to the initial condition so a single starting value sufficed.

### Statistics

All statistical comparisons were performed in MATLAB using a 2-sample Student’s *t*-test whose null hypothesis claimed there was no difference between the means of the two populations being tested. The populations in our analysis were compartmental and gradient ^23^Na signal values between tumor and normal tissue and between cohorts of different tumors. For ^1^H-DCE-MRI studies, the populations were different parameter values between different tumors. In all cases, a significance level of 0.05 was used.

## Supporting information

Theory and Supplemental Figures

## Data Availability

Data supporting the findings of this manuscript are available from the corresponding authors upon request.

## Acknowledgements

Research was supported by grants from the National Institute of Health awarded to F.H. (R01 EB-023366) and J.J.W. (T32 GM007205, Yale Medical Scientist Training Program).

## Author Contributions

M.H.K., D.C. and F.H. designed experiments. M.H.K., J.J.W. and J.M.M. conducted experiments and conducted data analysis. M.H.K., J.J.W. and S.K.M. prepared tumor cells. D.C. and F.H. supervised experiments and analysis. M.H.K., J.J.W., J.M.M. and F.H. evaluated results and wrote the manuscript.

