## Supplementary material for "Imaging the Transmembrane and Transendothelial Sodium Gradients in Gliomas": Theory and Supplemental Figures

**Supplementary: THEORY**

Complexation of Tm^3+^ with DOTP^8−^ stably(Sherry et al., 1996) forms TmDOTP^5^^−^ (**Figure 1(a)**), which attracts Na^+^ effectively due to its high negative charge. When intravenously injected, TmDOTP^5−^ resides in Na^+^-rich blood and extracellular compartments ([Na^+^]_b_ ≈ [Na^+^]_e_ ≈ 150 mM). Without loss of generality, the narrative below is with regard to Na^+^_e_, but also applies to Na^+^_b_. However, Na^+^_i_ is not affected by TmDOTP^5-^ (**Figure 1(b)**).

The total ^23^Na chemical shift depends on cumulative effects of diamagnetic and paramagnetic terms. The former has intramolecular and intermolecular contributions (e.g. ring currents, chemical exchange), but latter is affected by proximity of the nuclear spin to unpaired electrons(Abragam, 1961; Dwek, 1973). Electrostatic attraction forces Na^+^_e_ close to the paramagnetic Tm^3+^ core, whose unpaired electrons generate an intrinsic magnetic field **B_int_** augmenting the static magnetic field **B_0_**. This results in a shift of the ^23^Na resonance frequency from ω_0_ = γ_Na_**|B_0_|** to ω = γ_Na_**|B_0_+B_int_**| for Na^+^_e_, where γ_Na_=11.26 MHz/T and the chemical shift is δ≡(ω−ω_0_)/ω_0_ in units of parts per million (ppm).

Introducing paraCA*^n-^* creates a fast-exchange equilibrium between bound and unbound Na^+^_e_(Laszlo, 1978), thereby producing a single resonance detected between the two extremes. If the unbound and bound Na^+^_e_ pools have the chemical shifts δ*_u_* and δ*_b_* with respect to a reference frequency, then the observed ^23^Na peak has a chemical shift δ given by the weighted average of δ*_u_* and δ*_b_*:

$\delta=f_{b}\delta_{b}+f_{u}\delta_{u}$	**(1)**

where *f_b_* and *f_u_* are the respective fractions of Na^+^_e_ bound and unbound to TmDOTP^5−^ (*f_b_*+*f_u_*=1). The reference frequency is usually defined as that of natural (unbound) sodium (δ*_u_*~0). *In vivo*, the chemical shift caused by binding shows dependencies on temperature (*T*), pH, and [paraCA*^n-^*](Coman, Trubel, Rycyna, et al., 2009; Puckeridge et al., 2012):

$\delta(T,pH,[{paraCA}^{n-}])=\left( \frac{\partial\delta}{\partial T} \right)\Delta T+\left( \frac{\partial\delta}{\partial pH} \right)\Delta pH+\left( \frac{\partial\delta}{\partial[paraCA^{n-}]} \right)\Delta[{paraCA}^{n-}]$ **(2)**

These partial derivatives, or shiftability terms ($s_{f}\equiv\frac{\partial\delta}{\partial f}$), reflect the sensitivity of ^23^Na chemical shift to variations in their respective factors

$\delta(T,pH,[{paraCA}^{n-}])=s_{T}\Delta T+s_{pH}\Delta pH+s_{\left[ paraCA^{n-} \right]}\Delta[{paraCA}^{n-}]$	**(3)**

where $s_{T}, s_{pH}, and s_{[{paraCA}^{n-}]}$ are the respective *shiftabilities* for changes in temperature, pH and [paraCA*^n-^*] (**Figure 1(c)**). **Equation (3)** is based on shift changes for small variations to approximate linear limits of cation-anion binding models. *In vitro* tests reveal that [paraCA*^n-^*] has the greatest contribution to δ (**Figure 1(d,e)**) when considering realistic pH and/or temperature ranges experienced *in vivo* (**Figure 1(f,g)**). If we examine the ^23^Na chemical shift dependence on *relative* concentration [paraCA*^n-^*]/[Na^+^] as expected *in vivo* (i.e. ~2 mM vs. ~150 mM), then **Equation (3)** simplifies even further to

$\delta([paraCA^{n-}])\approx s_{[{paraCA}^{n-}]}\Delta[{paraCA}^{n-}]$	**(4)**

Note that ^23^Na-MRSI *shiftability* reflected by **Equation (4)** (in units of ppm shifted per mM) is analogous to the ^1^H MRI *relaxivity* (*r*) of a contrast agent (in units of rate enhanced per mM). Integrating the shifted peaks enables Na^+^ quantification under fully relaxed conditions. TmDOTP^5−^ induces a chemical shift in the Na^+^_b_ and Na^+^_e_ signals, with the former hypothesized to be shifted more than the latter based on [paraCA*^n-^*] in each compartment(Coman, Trubel, & Hyder, 2009; Coman, Trubel, Rycyna, et al., 2009). The integral of the shifted Na^+^_b_ peak reflects [Na^+^]_b_, and likewise the shifted Na^+^_e_ and unshifted Na^+^_i_ peaks, respectively, reflect [Na^+^]_e_ and [Na^+^]_i_.

**Supplementary: FIGURES**

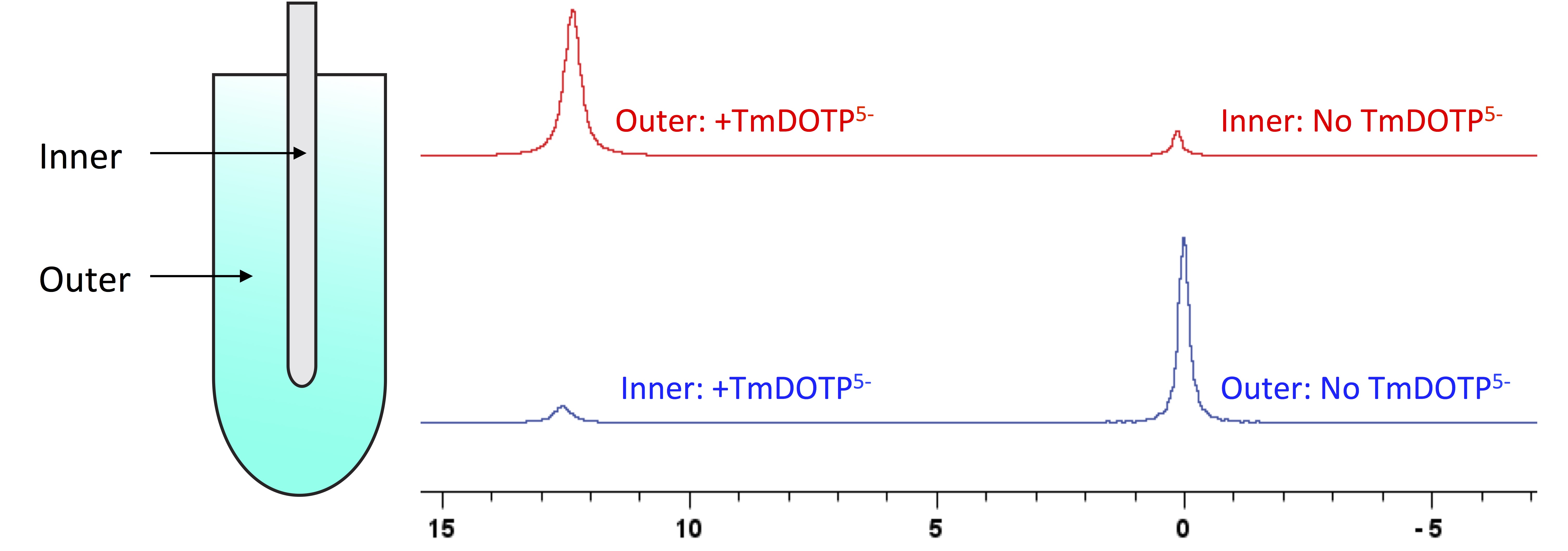

**Figure S1.** A diagram of the two-compartment coaxial tube setup used for *in vitro* experiments (left). The overall tube length was 7 inches, with a 5-mm body in which the second compartment (inner/outer diameter = 1.258/2.020 mm) was inserted. The length of the tip within the excitable volume was 50 mm. The outer-to-inner volume ratio was 8.6. A confirmatory study was conducted in which the inner compartment contained 150 mM NaCl and no TmDOTP^5−^ and the outer contained 150 mM NaCl with 5 mM TmDOTP^5−^ (right, red). The pH was set to 7.4, and the temperature in the spectrometer was 300 K. The outer compartment also contained D_2_O for frequency locking of the spectrometer. The resulting unshifted peak at 0 ppm and shifted peak at 12.5 ppm were consistent with reported apparent shiftability (*s*_app_) of 2.77 ppm/mM. Integration of the spectra yielded a shifted-to-unshifted integral ratio of 10, which reflects the ratio of moles between the two compartments. The contents of the compartments were switched (right, blue) and the resulting spectrum showed the peaks having interchanged positions and an inverted integral ratio, i.e. 0.1. Both compartments displayed broader peaks with TmDOTP^5−^ than without. This illustrates that TmDOTP^5−^ caused the observed shift effects and that only the difference in compartment volume was responsible for the size of the peak. See **Figure 1** for more example of representative data from this phantom setup.

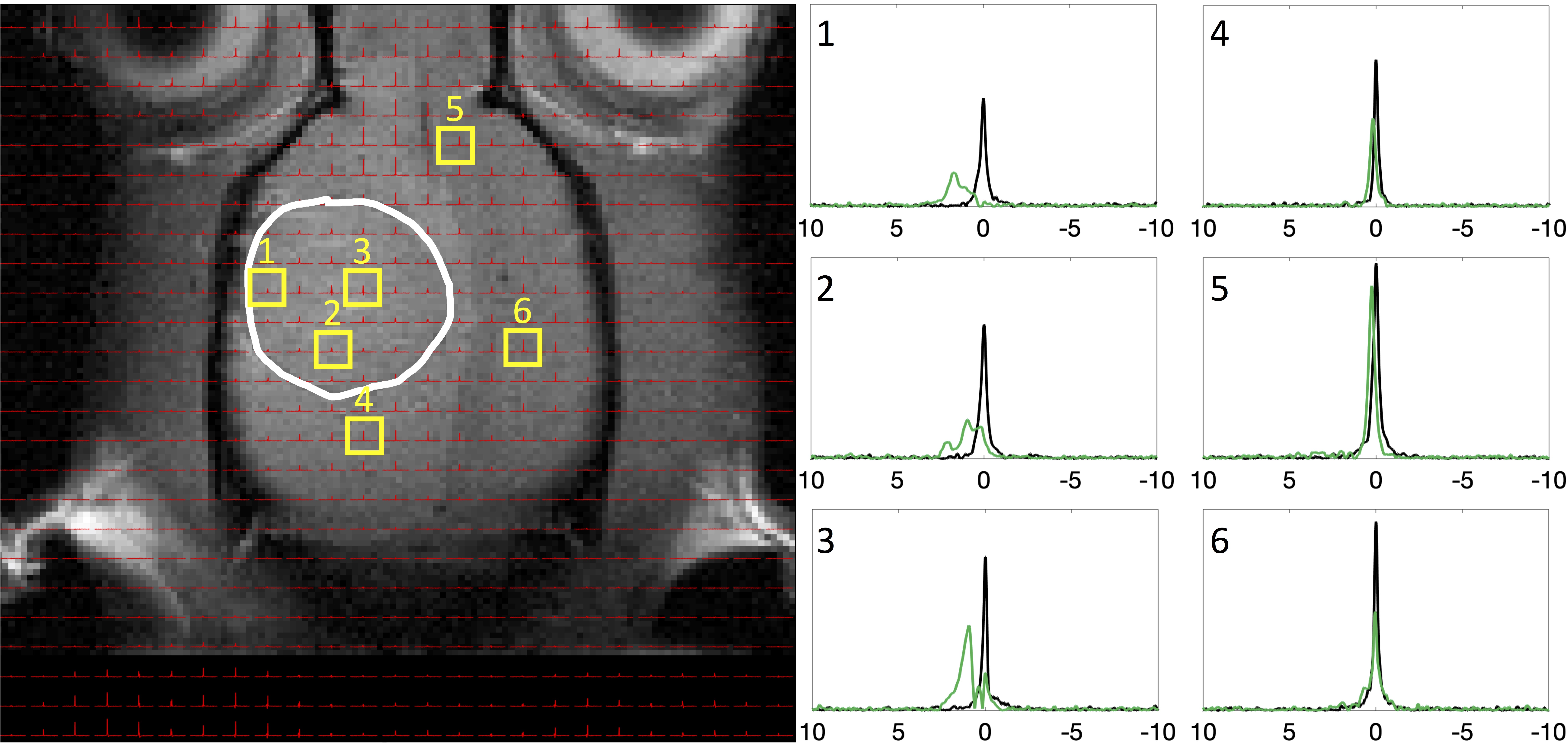

**Figure S2.** Demonstration of ^23^Na peak separation *in vivo* following TmDOTP^5−^ administration into a rat brain bearing a U251 tumor. ^1^H MRI of an axial slice displaying the anatomical tumor boundary (white outline). The ^23^Na-MRSI is overlaid on top of the MRI. Candidate voxels inside and outside the tumor are indicated (yellow boxes with numbers). Before delivery of the agent, a single ^23^Na peak was observed at 0 ppm in all voxels (black spectra), corresponding to total sodium (Na^+^_T_). Following TmDOTP^5−^ delivery, compartmental peak separation was achieved to varying extents throughout the brain (green spectra). Tumor voxels (1-3) exhibited a fair amount of peak separation due to the leaky blood-brain barrier (BBB), with blood sodium (Na^+^_b_ ) consistently around 2 ppm, extracellular sodium (Na^+^_e_ ) in the range 0.5-1 ppm, and the unshifted intracellular sodium (Na^+^_i_ ) at 0 ppm. Voxels outside the tumor (4-6) were slightly shifted in the positive direction, suggesting the paramagnetic effects of TmDOTP^5−^ reach the extracellular space even with limited extravasation. Similar spectroscopic patterns are observed throughout all voxels *in vivo*. See **Figure 2** for a slice above the present. All spectra were magnitude-corrected and line-broadened by 10 Hz.

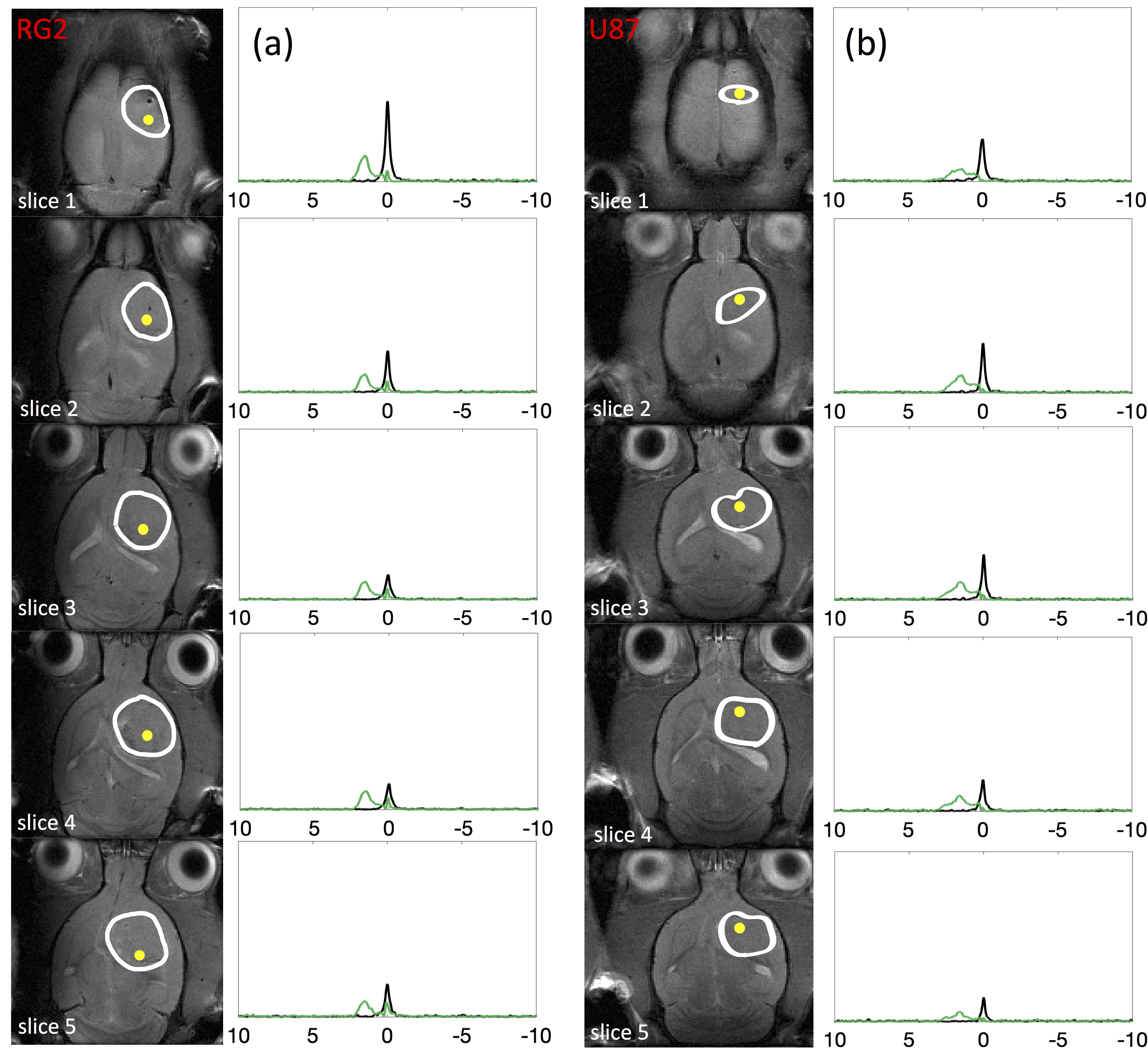

**Figure S3.** Comparison of ^23^Na peak separation in rats bearing an (a) RG2 and (b) U87 tumor. The tumor boundary is outlined in white, spectra acquired before and after TmDOTP^5−^ delivery are shown in black and green, respectively, and the yellow dots represent the same transverse coordinates across multiple slices. In both cases, “slice 1” is the most superficial axial slice with a visible tumor and slice thickness is 1 mm with no gap. The Na^+^_b_ peak was present at 2 ppm throughout the entire depth of the brain in both tumors, and was more shifted than Na^+^_e_, which was encountered around 0.5-1 ppm. The Na^+^_i_ peak was observable at 0 ppm. Similar spectroscopic patterns are observed throughout all voxels *in vivo*. See **Figure 3** for detailed comparison of voxels inside and outside these tumors. All spectra were magnitude-corrected and line-broadened by 10 Hz.

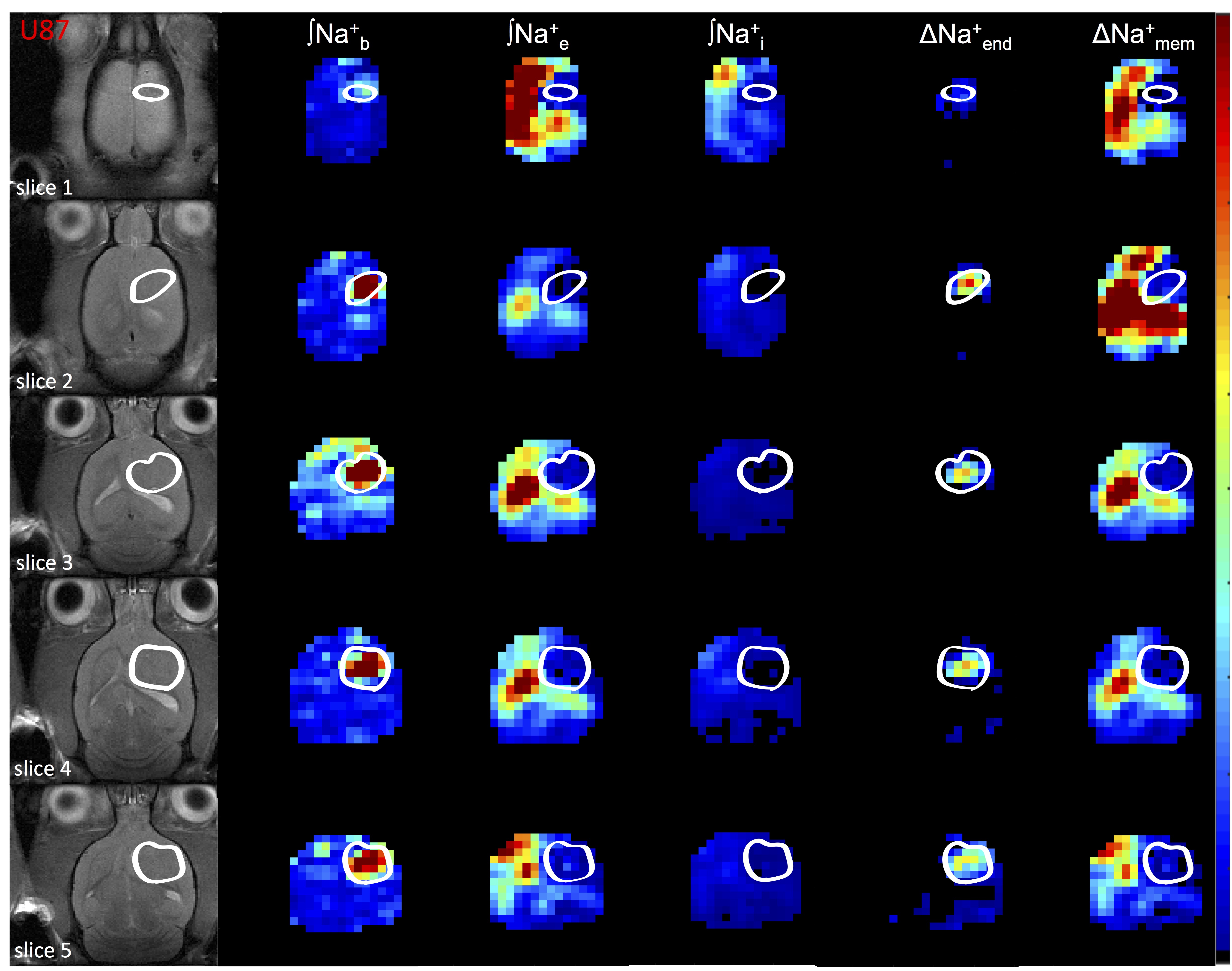

**Figure S4.** Spatial distributions of compartmentalized ^23^Na signals (Na^+^_b_, Na^+^_e_, Na^+^_i_) as well as transendothelial (ΔNa^+^_end_) and transmembrane (ΔNa^+^_mem_) gradients in an U87 tumor. The left column shows the tumor location (white outline) on the anatomical MRI (left). Since integral of ^23^Na peak in each compartment represents the [Na^+^], the respective three columns show the integral maps of Na^+^_b_, Na^+^_e_, and Na^+^_i_ from left to right (i.e., **∫**Na^+^_b_, **∫**Na^+^_e_, **∫**Na^+^_i_). The last two columns on the right show the ΔNa^+^_end_ = **∫**Na^+^_e_, **∫**Na^+^_i_ and ΔNa^+^_mem_ = **∫**Na^+^_e_-**∫**Na^+^_i_. The **∫**Na^+^_b_ map reveals low values in healthy tissue compared to tumor tissue, and within the tumor boundary high degree of heterogeneity is visible. The **∫**Na^+^_e_ map reveals low values in tumor and normal tissues, but within the tumor boundary small degree of heterogeneity is visible while ventricular voxels show very high values. The **∫**Na^+^_i_ map reveals low values ubiquitously except some ventricular voxels. The ΔNa^+^_end_ map reveals dramatically high values within the tumor only. The ΔNa^+^_end_ was driven primarily by an increase of **∫**Na^+^_b_ inside the tumor and which was more pronounced in superficial regions of the brain compared to deeper slices. The ΔNa^+^_mem_ map shows low values in tumor tissue compared to normal tissue, although ventricular voxels show very high values. The ΔNa^+^_mem_ is driven primarily by decreased **∫**Na^+^_e_ and thus shows similar level of heterogeneity as the **∫**Na^+^_e_ map. All maps use the same color scale and are relative. See **Figure 4** for an example for a RG2 tumor.

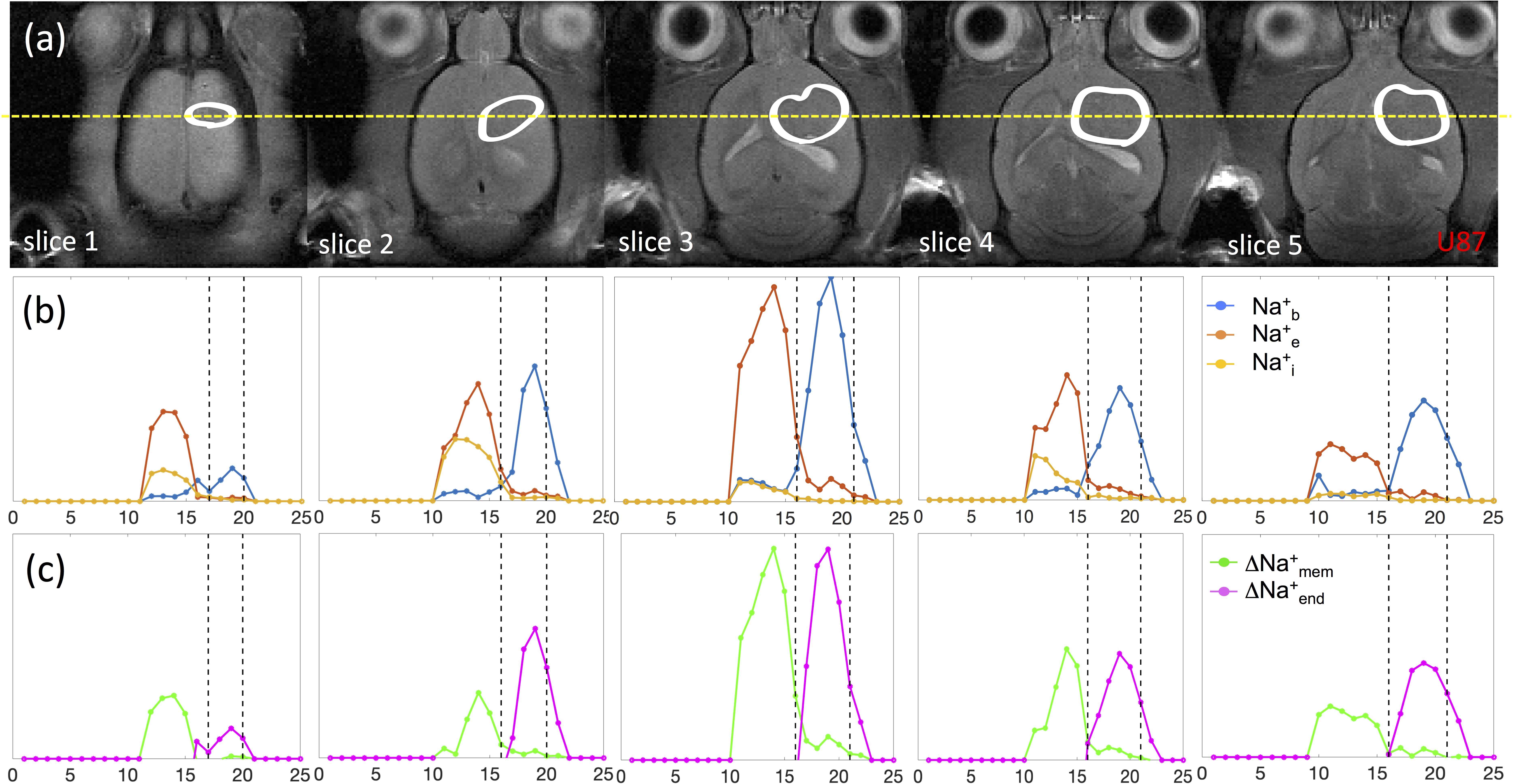

**Figure S5.** Coronal projections of compartmentalized ^23^Na signals (Na^+^_b_, Na^+^_e_, Na^+^_i_) as well as transendothelial (ΔNa^+^_end_) and transmembrane (ΔNa^+^_mem_) gradients in an U87 tumor. (a) Axial MRI indicating the tumor (white outline) across slices (same as **Figure S4**), where the yellow line indicates the position for a coronal projection. (b) Spatially varying ^23^Na signals for Na^+^_b_, Na^+^_e_, and Na^+^_i_ are shown with blue, orange, and yellow lines, respectively, where the vertical black lines indicate the tumor boundary. The Na^+^_b_ signal (blue) is clearly elevated in the tumor, and most elevated in slice 3 (or medially). This pattern is similar for Na^+^_e_ signal (orange) but the elevation is outside the tumor. Thus behavior of Na^+^_b_ signal (blue) is inversely related to Na^+^_e_ signal (orange). Both intratumoral Na^+^_b_ signal (blue) and peritumoral Na^+^_e_ signal (orange) are highest in slices 3. The Na^+^_i_ signal (yellow) varies slightly across slices. It is highest in slices 1-2, and negligible in slice 3-5. (c) Behaviors of ∆Na^+^_mem_ signal (green) and ∆Na^+^_end_ signal (magenta) closely mimic patterns of Na^+^_e_ and Na^+^_b_ signals, respectively, indicating that each of those Na^+^ compartments is the primary driver of the respective Na^+^ gradient. See **Figure 5** for a similar example for a RG2 tumor.

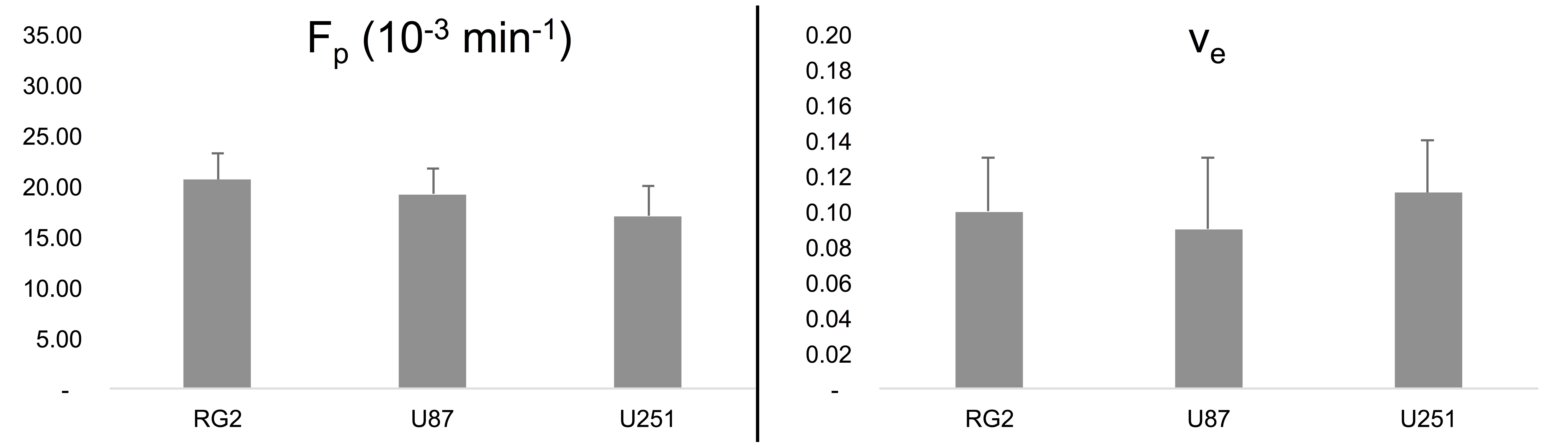

**Figure S6:** Comparisons of *F_p_* and *v_e_* ^1^H DCE-MRI parameters between RG2, U87 and U251 tumors. For all tumor types, mean *F_p_* values (left) closely followed patterns for *K^trans^* (see **Figure 6(c)**), but there was only a marginally significant difference between tumor types. There was also no significant difference between *v_e_* values (right) for the different tumor types. These values, however, are smaller than the observed *v_p_* values (see **Figure 6(c)**), which suggests that the vascular portion of all tumor types studied dominates over the interstitial space. See **Figure S7** for exemplary maps of ^1^H-DCE-MRI parameters for animals from each tumor type.

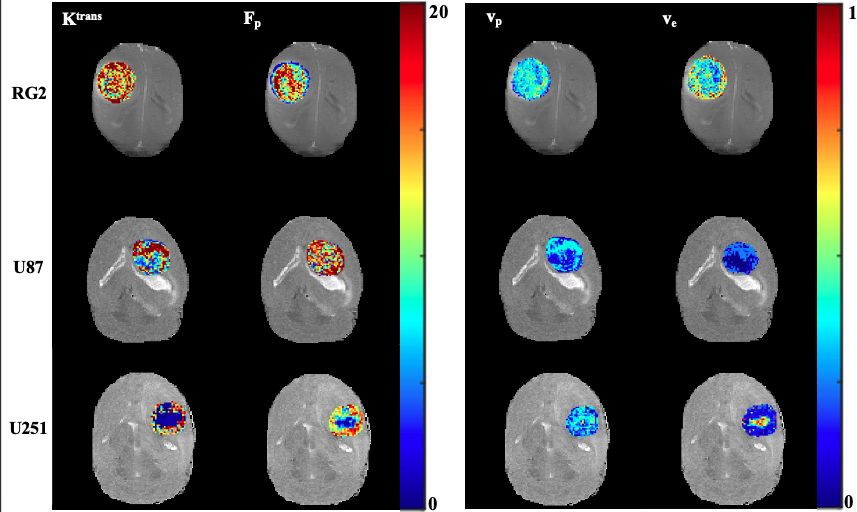

**Figure S7.** Exemplary maps of ^1^H-DCE-MRI parameters from representative animals from each tumor type. ^1^H-DCE-MRI parameters were calculated only for the tumor, and overlaid on *T_1_­*-weighted anatomical images. Values for *K^trans^* and *F_p_* are reported in 10^-3^ min^-1^. As measures of vascularity, these values reveal heterogeneity throughout all three tumor types, but overall significantly larger values are present for RG2 and U87 relative to U251 (see **Figure 6(c)**). Small regions with low *K^trans^* and *F_p_* near the center of the U251 tumor suggest areas of low perfusion and are indicative of a necrotic core, which U251 tumors have been shown to have. Furthermore, *v_p_* values tend to be larger throughout the tumor compared to *v_e_* which suggests the large presence of vasculature within tumor tissue. The necrotic core in U251, defined by the low-*F_p_* region within the tumor, shows a corresponding region of elevated *v_e_* suggesting the lack of vasculature in areas with low perfusion. These findings help to confirm the elevated Na^+^_b_ signals observed in ^23^Na-MRSI are consequences of enhanced vascularity. See **Figure 6(c)** for the distribution of *K^trans^* and *v_p_* values, and **Figure S6** for the distribution of *F_p_* and *v_e_* values.

Abragam, A. (1961). *The principles of nuclear magnetism*. Clarendon Press.

Coman, D., Trubel, H. K., & Hyder, F. (2009). Brain temperature by Biosensor Imaging of Redundant Deviation in Shifts (BIRDS): comparison between TmDOTP 5âand TmDOTMA â. *NMR in Biomedicine, 29*(2), n/a-n/a. <https://doi.org/10.1002/nbm.1461>

Coman, D., Trubel, H. K., Rycyna, R. E., & Hyder, F. (2009, Feb). Brain temperature and pH measured by 1H chemical shift imaging of a thulium agent. *NMR in Biomedicine, 22*(2), 229-239. <https://doi.org/10.1002/nbm.1312>

Dwek, R. A. (1973). *Nuclear magnetic resonance (N.M.R.) in biochemistry : applications to enzyme systems*. Clarendon Press.

Laszlo, P. (1978, Apr). Sodium-23 Nuclear Magnetic Resonance Spectroscopy. *Angewandte Chemie International Edition in English, 17*(4), 254-266. <https://doi.org/10.1002/anie.197802541>

Puckeridge, M., Chapman, B. E., Conigrave, A. D., & Kuchel, P. W. (2012, Oct). Quantitative model of NMR chemical shifts of 23Na+ induced by TmDOTP: Applications in studies of Na+ transport in human erythrocytes. *Journal of Inorganic Biochemistry, 115*, 211-219. <https://doi.org/10.1016/j.jinorgbio.2012.03.009>

Sherry, A. D., Ren, J., Huskens, J., Brücher, E., Tóth, É., Geraldes, C. F. C. G., Castro, M. M. C. A., & Cacheris, W. P. (1996, Jan). Characterization of Lanthanide(III) DOTP Complexes:  Thermodynamics, Protonation, and Coordination to Alkali Metal Ions. *Inorganic chemistry, 35*(16), 4604-4612. <https://doi.org/10.1021/ic9600590>
